# Non-nucleosomal (CENP-A/H4)_2_ - DNA complexes as a possible platform for centromere organization

**DOI:** 10.1101/2024.12.31.630874

**Authors:** Ahmad Ali-Ahmad, Mira Mors, Manuel Carrer, Xinmeng Li, Silvija Bilokapić, Mario Halić, Michele Cascella, Nikolina Sekulić

## Abstract

The centromere is a part of the chromosome that is essential for the even segregation of duplicated chromosomes during cell division. It is epigenetically defined by the presence of the histone H3 variant CENP-A. CENP-A associates specifically with a group of 16 proteins that form the centromere-associated network of proteins (CCAN). In mitosis, the kinetochore forms on the CCAN to connect the duplicated chromosomes to the microtubules protruding from the cell poles. Previous studies have shown that CENP-A replaces H3 in nucleosomes, and recently the structures of CENP-A-containing nucleosomes in complex with CCANs have been revealed, but they show only a limited interaction between CCANs and CENP-A. Here, we report the cryoEM structure of 2x(CENP-A/H4)_2_-di-tetramers assembled on DNA in the absence of H2A/H2B histone dimer and speculate how (CENP-A/H4)_2_-tetramers and -di-tetramers might serve as a platform for CCAN organization.

**Main points:**

- CENP-A/H4 in the absence of H2A/H2B dimers associate on DNA to form di-tetrasomes
- CENP-A/H4 di-tetrasomes are stable but highly flexible structures resembling nucleosomes
- CENP-A/H4 tetrasomes bind CENP-N/L and can serve as platform for centromere organisation

## Introduction

The centromere plays a crucial role in the correct distribution of genetic material during cell division by ensuring that the duplicated chromosomes are divided evenly between two new cells (1–5). The centromere is epigenetically defined by the presence of an H3 histone variant called CENP-A (reviewed in (6)). CENP-A is instrumental in the organization of the constitutive centromere-associated network (CCAN) of 16 proteins. Despite major advances in the obtaining an atomic resolution of CENP-A nucleosomes (7, 8) and CCANs (9, 10) and even cryoEM structures of human and yeast CENP-A nucleosomes in complex with CCANs (11–13), it is still not clear which CENP-A features control CCAN organization in CENP-A-enriched parts of chromatin. Furthermore, it has been proposed that CCAN structure and the nature of its interaction with CENP-A changes dynamically during the cell cycle (1, 14–16).

There are several indications that only two CCAN components, CENP-C and CENP-N, can interact directly with CENP-A and that the remaining proteins form subcomplexes on this structure ((17) and also reviewed in (18)). The two segments on the human CENP-C (CENP-C^426-537^ and CENP-C^737-359^) recognize and bind the nucleosome CENP-A by interlocking with its C-terminal tail, which is exposed on the surface of the nucleosome and acidic patch on histone H2A that is also exposed on the surface of the nucleosome. The interaction between CENP-C and CENP-A nucleosome has been visualized in several studies, including the CENP-A/CCAN structures of yeast and human complexes (8, 11–13, 16, 19, 20). On the other hand, CENP-N was found to interact with the L1 loop of CENP-A, which has an RG insertion (compared to the H3 histone) (21). The RG loop is also exposed on the surface of the CENP-A nucleosome and several cryoEM structures have captured the interaction of the N-terminal portion of CENP-N with the CENP-A^RG-loop^ and DNA on the CENP-A nucleosome (16, 22–24). However, when full-length CENP-N is presented together with the binding partner CENP-L in the context of CCANs, the CENP-N/L complex does not bind the CENP-A^RG-loop^ on CENP-A nucleosomes, but rather binds only DNA protruding from the nucleosome and far away from the histone core (11–13). This leaves open the question in which context CENP-A and CENP-N interact directly via the CENP-A^RG-loop^ and whether it is possible that either CENP-A or CCAN have an alternative organization in chromatin that allows direct interaction between CENP-A and CENP-N during the cell cycle.

Previous high-resolution structures of CENP-A show either (CENP-A/H4)_2_ tetramers without DNA (25) or nucleosomes (multiple structures, in isolation or in complex with other proteins), suggesting that these are the most stable complexes of CENP-A. Several proposals for CENP-A-containing particles have been made in the past (reviewed in (26)) including a CENP-A “hemisome” structure in which only one copy of each histone (CENP-A, H4, H2A and H2B) is present. However, this particle has not been stabilized *in vitro* for high-resolution studies, nor has it been observed in cells (27). To reconcile the cryoEM structure of CCANs with previous observations of CENP-A/CENP-N interaction, Musacchio’s lab proposed a “hemisome” in which CENP-T/CENP-W replaces H2A/H2B, but the high-resolution structure of the particle was never reported (9).

Here, we show that (CENP-A/H4)_2_ can form a di-tetrasome structure on DNA *in vitro* in the absence of H2A/H2B histones. We report two different conformations captured by cryoEM that we call open and closed. The MD analysis finds that these structures are stable but highly dynamic and we provide evidence that 2x(CENP-A/H4)_2_ di-tetrasomes can form a unique chromatin structure *in vitro*. We also show that (CENP-A/H4)_2_ di-tetrasome can bind CENP-C albeit with lower affinity than CENP-A nucleosome and that (CENP-A/H4)_2_ tetrasome can bind CENP-N/L complex. We discuss the implications of our findings on the possible interactions between CENP-A and CCANs.

## Material and Methods

### Protein cloning and purification

#### (H3/H4)_2_ and (CENP-A/H4)_2_

were expressed as heterotetramers. (H3/His-H4)_2_ and (His-CENP-A/H4)_2_ were each cloned into pst44 plasmids and bicistronically expressed in *E. coli* Rosetta2 cells overnight at 18°C. Harvested cells were resuspended in lysis buffer and sonicated. The supernatant was injected onto a 5 ml HisTrap HP (Cytiva), and (H3/His-H4)_2_ or (His-CENP-A/H4)_2_ were eluted with buffer containing 20 mM Tris pH 7.5, 2 M NaCl, 5% glycerol, 5 mM BME (β-mercaptoethanol) and 300 mM imidazole. The His tag was cleaved overnight with in-house prepared TEV protease, and sample was subjected to a final cation exchange chromatography step (Hitrap SP HP 5ml; Cytiva). Elution was performed in steps with the final buffer containing 20 mM Tris pH 7.5, 2 M NaCl and 2 mM DTT.

#### CENP-N^1-240^

CENP-N^1-240^ with a His tag on the C-terminus was expressed overnight at 18 °C in *E. coli* pLysS. Pelleted cells were dissolved in 50 mM NaPO_4_ pH 8, 300 mM NaCl, 10 mM imidazole, 0.1 % Tween-20, and lysed by sonication. CENP-N^1-240^ was purified using a 5 mL HisTrap FF column, and eluted with 50 mM NaPO_4_ pH 7,5, 500 mM NaCl, 300 mM imidazole, 20% glycerol. The eluted protein was further purified on a Superdex S200 16/600 size exclusion column preequilibrated with 50 mM NaPO_4_ pH 7,5, 500 mM NaCl, 20% glycerol.

#### CENP-N/L

The human CENP-N/His-CENP-L complex was expressed in Sf9 cells for 3 days at 27.5°C (Lund protein production platform) as described before (28). Harvested cells were resuspended in buffer containing 20 mM Tris pH 7.5, 0.3 M NaCl, 5% glycerol, 5 mM BME, 0.05 % Tween 20, EDTA-free anti-protease cocktails (Sigma) and sample was sonicated. The supernatant was injected onto a 5 ml HisTrap HP (Cytiva), and CENP-N/L complex was coeluted using 20 mM Tris pH 7.5, 0.3 M NaCl, 5 % glycerol, 5 mM BME and 300 mM Imidazole. A further size exclusion chromatography purification step was performed using S200 column (Cytiva) pre-equilibrated in 20 mM HEPES 7.5, 300 mM NaCl, 2 mM DTT.

#### CENP-H/I/K/M

His-CENP-H/K/his-CENP-I^57-C^/M heterotetramer was cloned into the pst44 plasmid and coexpressed from tetracystronic mRNA in *E. coli* Rosetta2 cells overnight at 18°C. Harvested cells were resuspended in 20 mM Tris pH 7.5, 0.5 M NaCl, 5% glycerol, 5 mM BME, and EDTA-free anti-protease cocktail (Sigma) and lysed by sonication. The supernatant was injected onto a 5 ml HisTrap HP (Cytiva), and the eluted complex was further purified using anion exchange chromatography column (Hitrap Q HP 5ml; Cytiva). Pure His-CENP-H/K/his-CENP-I/M was eluted using a salt gradient with the final buffer composition 20 mM Tris pH 7.5, 1M NaCl, 2mM DTT.

#### CENP-C

CENP-C^426-537^ was expressed in E. coli pLysS and purified as previously described in (29). Briefly, GST-tagged CENP-C^426-537^ was purified on a glutathione column (GSTrap™ HP Cytiva). GST was subsequently cleaved overnight by PreScission protease and separated from CENP-C using cation exchange chromatography (Hitrap SP HP 5ml; Cytiva).

### DNA preparation

145 bp and 200 bp 601 super-positioning DNAs and 147 bp α-satellite DNA were purified as described in (29, 30). Briefly, HB101 cells transformed with pUC57 plasmids containing 6 × 147 bp α-satellite DNA or 8 × 145 bp 601 or 19x 200bp 601 super-positioning DNA (gift from Ben Black, UPenn) were grown at 37C overnight, and DNA was extracted with phenol/chloroform. Plasmids were digested using restriction enzymes and further purified using anion-exchange chromatography (resource Q 6ml column, Cytiva).

601 (145 bp):

ATCAGAATCCCGGTGCCGAGGCCGCTCAATTGGTCGTAGACAGCTCTAGCACCGCTTAAACGCACGTACGCGCTGTCCCCCGCGTTTTAACCGCCAAGGGGATTACTCCCTAGTCTCCAGGCACGTGTCAGATATATACATCGAT

601 (200 bp):

TATGTGATGGACCCTATACGCGGCCGCCCTGGAGAATCCCGGTGCCGAGGCCGCTCAATTGGTCGTAGCAAGCTCTAGCACCGCTTAAACGCACGTACGCGCTGTCCCCCGCGTTTTAACCGCCAAGGGGATTACTCCCTAGTCTCCAGGCACGTGTCAGATATATACATCCTGTGCATGTATTGAACAGCGACTCGGGT

α-satellite (147 bp):

ATCAAATATCCACCTGCAGATTCTACCAAAAGTGTATTTGGAAACTGCTCCATCAAAAGGCATGTTCAGCTCTGTGAGTGAAACTCCATCATCACAAAGAATATTCTGAGAATGCTTCCGTTTGCCTTTTATATGAACTTCCTCGAT

di-α-satellite (342bp):

CTGAGGCCTGTGGTAGTAAAGGAAAGAACTTCATATAAAAACTAGACGGTAGCACCCTCAGAAAATTCTTTGTGACGATGGAGTTTAACTCAGAGAGCTGAACATTCGTTATGATGGAGCAGTTTCCAAACACACGTTTTGTAGAATCTGCAAGGGGATATTTGGACCTTCCGGAGGATTTCGTTGGAAACGGGATCAACTTCCCATAACTGAACGGAAGCAAACTCAGAACATTCTTTGTGATGTTTGTATTCAACTCACAGAGTTGAACCTTCCTTTGATAGTTCAGGTTTGCAACACCCTTGTAGTAGAATCTGCAAGTGTATATTTTGACCACTTTGG

### Di-tetrasome and poly-tetrasome complexes assembly

Di-tetrasome and poly-tetrasome complexes were assembled using salt gradient dialysis. (CENP-A/H4)_2_ or (H3/H4)_2_ hetero-tetramers were mixed with 601 (145 bp) or α-satellite (147 bp) DNA sequences in a molar ratio of 2:1 for di-tetrasome, and with di-α-satellite (342bp) DNA sequence in a molar ratio of 5:1 for poly-tetrasome in high salt buffer (20 mM Tris pH7.5, 2M NaCl, 2mM DTT). Gradient dialysis to low salt buffer (10 mM Tris pH 7.5, 100 mM NaCl, 2 mM DTT) was performed overnight at a flow rate of 1.5 ml/min using a dual-channel peristaltic pump. The next day, complexes were dialyzed against 10 mM MOPS 7.5, 100 mM NaCl, 2 mM DTT buffer and stored at 4°C, if necessary. complexes quality was checked using a 5% native PAGE gel.

### Binding assays

1 μM of 2x(CENP-A/H4)_2_ or 2x(H3/H4)_2_ di-tetrasome assembled on 601 (145 bp or 200bp) DNA or α-satellite DNA were mixed with various amounts of CENP-C^426-537^, CENP-N^1-240^, CENP-N/L, or CENP-H/I^57-C^/K/M and incubated for 1 h on ice. Complex formation was monitored using a 5% native PAGE gel.

### MNase experiments

0.5 μg of nucleosome, di-tetrasome or tetrasome complexes (based on DNA concentration) were incubated with 1 Kunitz unit of micrococcal nuclease (NEB) in a buffer containing 10 mM Tris HCl pH 7.5, 3 mM CaCl2, and 1 mM DTT at room temperature. Reactions were quenched at different time points (2, 5, 8, 10, and 15 min) by adding 250 μl PB buffer (Qiagen QIAquick PCR Purification Kit) supplemented with 10 mM EGTA. DNA from each sample was purified using the QIAquick PCR Purification Kit, and the extent of DNA digestion was quantified using the 2100 Bioanalyzer (Agilent). All experiments were performed in duplicate.

### Cryo-EM grids preparation, data acquisition and processing

The di-tetrasome complexes were cross-linked with 0.05% glutaraldehyde for 40 min on ice. 3 μl of the crosslinked 2x(H3/H4)_2_ di-tetrasome complex and uncrosslinked poly-tetrasome complex at a concentration of 0.8-1mg/ml was applied to freshly glow-discharged Quantifoil R2/1 300 mesh grids. Crosslinked 2x(CA/H4)_2_ di-tetrasome was applied on graphene oxide coated grids at a concentration of 0.1 mg/ml. All Grids were blotted for 5 seconds and frozen in liquid ethane using a FEI Vitrobot automatic plunge freezer. Humidity in the chamber was maintained at 100%.

2x(H3/H4)_2_ and 2x(CENP-A/H4)_2_ di-tetrasome datasets were acquired using the Titan Krios electron microscope (Thermo Fisher Scientific) at 300 kV (cryo-EM facility at UCEM, Umeå university, Sweden). Both datasets, 14209 movies for 2x(H3/H4)_2_ di-tetrasome particles and 12627 for 2x(CENP-A/H4)_2_, were acquired using a Gatan Summit K2 electron detector at a magnification of 165k and a pixel size of 0.82 Å. The total electron exposure of 57.5 e-/Å2 was distributed over 40 frames. (CENP-A/H4)_2_ poly-tetraseome dataset was acquired at UCEM cryo-EM Facility using the Titan Krios electron microscope (Thermo Fisher Scientific) at 300 kV. 8140 movies were acquired using Falcon4i electron detector at a magnification of 165k and a pixel size of 0.704 Å. The total electron exposure of 50 e-/Å2 was distributed over 828 frames.

Data were acquired using EPU (Thermo Fisher Scientific) automated data acquisition software with AFIS. The defocus range was from −1 to −3.8 μm with a step size of 0.3 μm.

Movie frames were aligned using patch motion in CryoSPARC 3.1 (31). CTF was estimated in a patch manner and several hundreds of particles were manually picked. The resulting useful particles were then used for automatic particle picking using Topaz (32). The 2D class averages were generated in CryoSPARC 3.1. Inconsistent class averages were removed from further data analysis. The 3D classifications and refinements were subsequently done in CryoSPARC 3.1 (Supplementary Figure 1). The initial reference was generated using ab initio and was filtered to 30 Å, and C1 symmetry was applied during homogenous refinements. Particles were split into two datasets and refined independently, and the resolution was determined using the 0.143 cut-off. Local resolution was determined with local resolution estimation. All maps were filtered to local resolution using CryoSPARC 3.1 with a B-factor determined in the refinement step.

Statistics on cryoEM maps are summarized in Table S1 and S2.

### Model building

The model was built in Coot (33) and refined using Phenix real_space_refine (34). Figures are prepared with Chimera (35).

### Molecular Dynamics

#### System setup

Molecular mechanics models for the 2x(H3/H4)_2_ or 2x(CEN-P/H4)_2_ di-tetrasomes were built starting from the experimental cryo-EM structures introduced in this work. As the experimental resolution allows for characterization of only the core protein regions of the complexes, models of the complete histones were created reconstructing the missing residues with the MODELLER package (36). Missing DNA ends not wrapped to the histones were added as canonical double helix moieties, using the web 3DNA server (37). For comparison, we also simulated conventional centromeric (pdb id: 6O1D) (38), and non-centromeric nucleosomes (pdb id: 3LZ0) (39) on 601 super positioning sequence (40), using 3D models from, respectively, cryo-EM data (3.4 Å resolution), and x-ray diffraction (2.5 Å resolution) as starting coordinates. All missing hydrogen were added using geometric restraints. The protonation states of all titratable residues were determined at pH 7, considering possible pKa shifts (41, 42). All the systems were solvated by TIP3P waters (43) setting the initial edge of the cubic periodic box to a length of 20 nm. Sodium and chloride ions were added to the system to achieve charge neutralization, and a saline concentration of 150 mM, mimicking physiological ionic strength. The resulting systems contained roughly 0.8 million of atoms (Table S3). Interaction energy terms were described using the Amber force field (44, 45), electrostatic interactions were computed using particle mesh Ewald (46), Lennard-Jones (LJ) potentials mimicking van der Waals interaction were approximated by cut-off scheme. A Cut-off radius of 1.2 nm was used for both the LJ and the short-range Electrostatics terms, updating the pair list every 20 molecular dynamics (MD) steps using a Verlet cut-off scheme (47). All simulations were run for 900 ns.

#### Simulation parameters

All the systems underwent several rounds of steepest descent energy minimization (maximum 10 000 cycles), and thermal equilibration by simulated annealing in the NPT to reach the target temperature of 300 K, targeted by the canonical velocity rescaling algorithm (48), using at coupling constant τ_T_ = 1.0 ps. Constant pressure of 1.0 Bar was achieved coupling MD simulations to stochastic cell rescaling barostat (49), using a coupling constant τ_p_ = 2.0 ps. The timestep for all the simulations was set to Δt = 2 fs, using the leap-frog integrator. The LINCS algorithm (50) was used to constrain all the bonds involving hydrogen atoms to their equilibrium distance. All simulations were performed using the GROMACS software (51).

#### Minimization of the AlphaFold models

After obtaining tetrasome/CCAN complex models using Alphafold, the protonation states of all titratable amino acids at neutral pH were determined by calculating their pKa values with PDB2PQR (52). The structures were then energy-minimized using Gromacs software (53) and the AMBER99bsc1 force field parameters (44, 45). Van der Waals interactions were calculated with a 1.0 nm cutoff distance, while electrostatic interactions were calculated using the Particle Mesh Ewald (PME) method (54). The steepest descent energy minimization was performed for up to 5000 steps, aiming for a convergence threshold with a force constant of 1000 kJ/(mol nm). Hydrogen bonds were constrained by the LINCS algorithm.

#### Measuring angle and distance for gyre opening

The gyre opening distance and angles were defined considering the base-pair of the wrapped DNA at the dyad location (base-pair residues numbered “0” in the 6O1D and di-tetrasome PDBs; base-pair residues numbered “73” in the 3LZ0 structure), and the base-pairs at the opposite location on the two DNA arms (± 39 in the DNA sequence).

The gyre distance and angles were defined as the distance between the N1 atoms of the purine nucleobase at ± 39 positions from the dyad, while the gyre angle was defined as the angle formed by the N1 atoms of the purine nucleobases at dyad-39, dyad, dyad+39 positions.

Measured angles and distances are summarized in Table S4.

## Results

### Two (CENP-A/H4)_2_ tetramers assemble into 2x(CENP-A/H4)_2_ di-tetrasome on DNA

We incubated (CENP-A/H4)_2_ tetramers or (H3/H4)_2_ tetramers with 147 bp DNA in 2:1 ratio in the absence of H2A/H2B dimer (Figure 1A) and we observed formation of two types of particles with different mobility on the native gel (Figure S1A). The particle corresponding to slower band on gel is tetrasome while the one that travels faster and almost the same as CENP-A nucleosome is 2x(CENP-A/H4)_2_ di-tetrasome (Figure S1A). This is similar to recent observation of 2x(H3/H4)_2_ di-tetrasomes by (55). Indeed, we could assemble di-tetrasomes with both (H3/H4)_2_ and (CENP-A/H4)_2_ tetramers on super-positing and on natural centromeric alpha-satellite DNA, albeit 2x(CENP-A/H4)_2_ di-tetrasomes consistently assemble with lower efficiency (Figure S1B). Next, we wondered if di-tetrasomes would protect wrapped DNA in the same way as nucleosomes, so we performed MNase digestion of assembled particles (Figure S1C). Not entirely surprisingly, nucleosomes wrap and protect DNA much more profoundly then di-tetrasomes, but di-tetrasomes nevertheless show some DNA protection that is less pronounced for 2x(CENP-A/H4)_2_ di-tetrasomes comparing to their H3-containing counterparts. To better understand these structures, we obtained the cryo-EM maps of 2x(CENP-A/H4)_2_ di-tetrasomes to 4.1 Å (Figure 1B and S2; Table S1) and 2x(H3/H4)_2_ di-tetrasomes to 3.75 Å resolution (Figure S3; Table S1).

**Figure 1:**
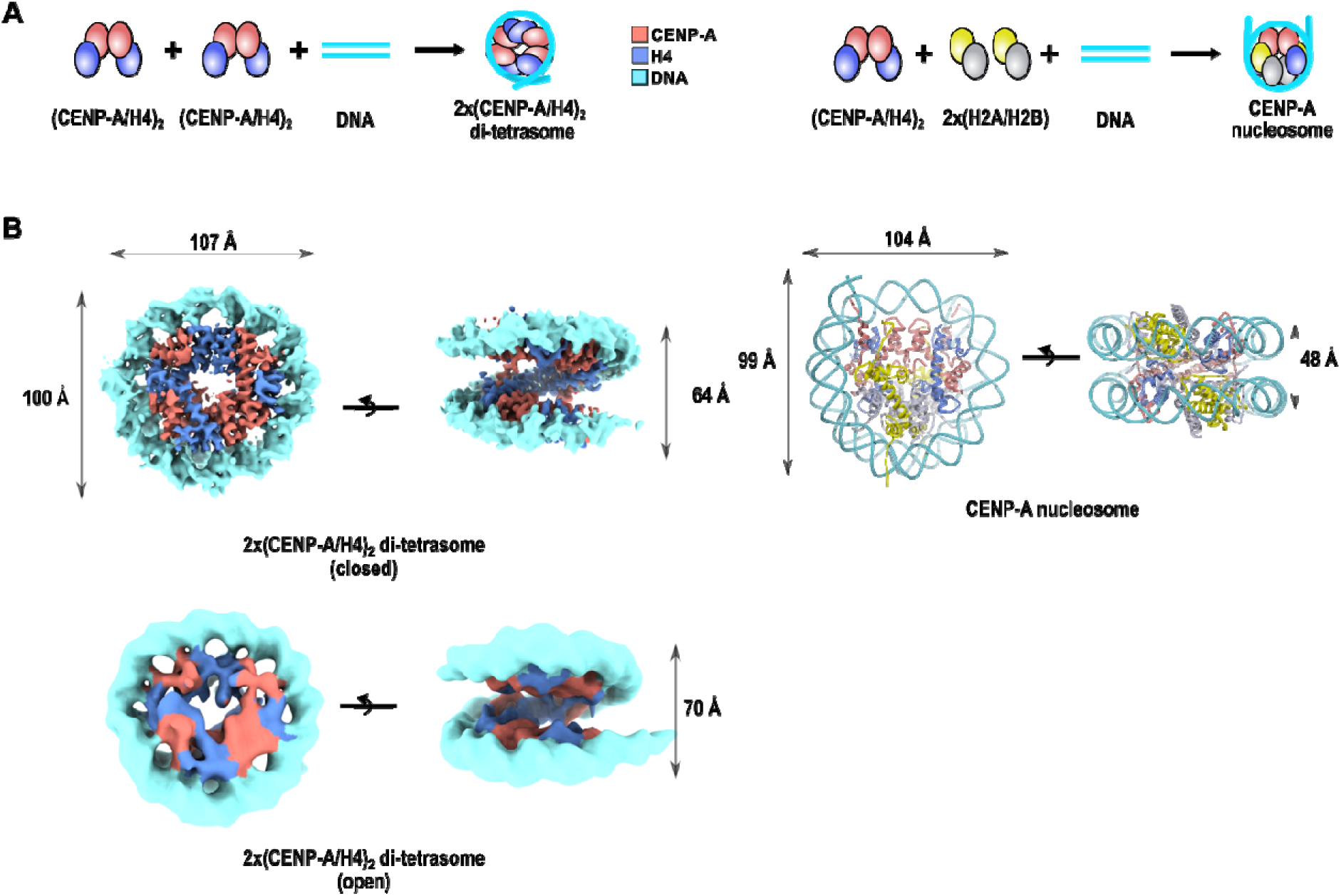
Two (CENP-A/H4)_2_ tetramers can assemble on DNA to form 2x(CENP-A/H4)_2_ di-tetrasomes. **A.** Schematic representation of the assembly of 2x(CENP-A/H4)_2_ di-tetrasomes (left) and CENP-A nucleosomes (right) for comparison. CENP-A is red; H4 is blue; H2A is yellow; H2B is gray; and 145 bp DNA is cyan. **B.** CryoEM map of the 2x(CENP-A/H4)_2_-di tetrasome (left) in open and closed conformation with measurements compared to the model CENP-A nucleosome (pdb ID: 6SE0) on the right. Histones and DNA are colored as in A. Note that the 2x(CENP-A/H4)_2_ di-tetrasome four-helix bundle at the dyad is formed by H4-H4 interactions.

### 2x(CENP-A/H4)_2_ di-tetrasome forms a flexible nucleosome-like particle

2x(CENP-A/H4)_2_ di-tetrasome have the same overall structure as 2x(H3/H4)_2_ di-tetrasome that was recently described (55). To be able to compare the structures directly, we have obtained the cryoEM structure of both 2x(H3/H4)_2_ and 2x(CENP-A/H4)_2_ di-tetrasomes in the same conditions. Analysis of cryoEM data indicates that di-tetrasomes are much more flexible (multiple conformations) than nucleosomes, which is also in agreement with higher susceptibility to MNase digestion (Figure S1C). For each of the di-tetrasome particles, we refined two different classes (open and closed) representing the most populated conformations (Figure 1B, S2B and S3D). 2x(CENP-A/H4)_2_ di-tetrasome have two (CENP-A/H4)_2_ tetramers accommodated on 147 bp DNA, so in difference to nucleosomes that has CENP-A/CENP-A four-helix bundle at the dyad, di-tetrasome have H4/H4 four-helix bundle at the dyad and two CENP-A/CENP-A four-helix bundles are on the superhelical location 3 (SHL3) and −3 (SHL-3) (Figure 2A-C). Also immediately noticeable is a greater separation of DNA gyres opposite of the dyad (Figure 2D). The detailed structural analysis reveals several differences between nucleosomes and di-tetrasomes that result in decreased compaction of the later.

**Figure 2.**
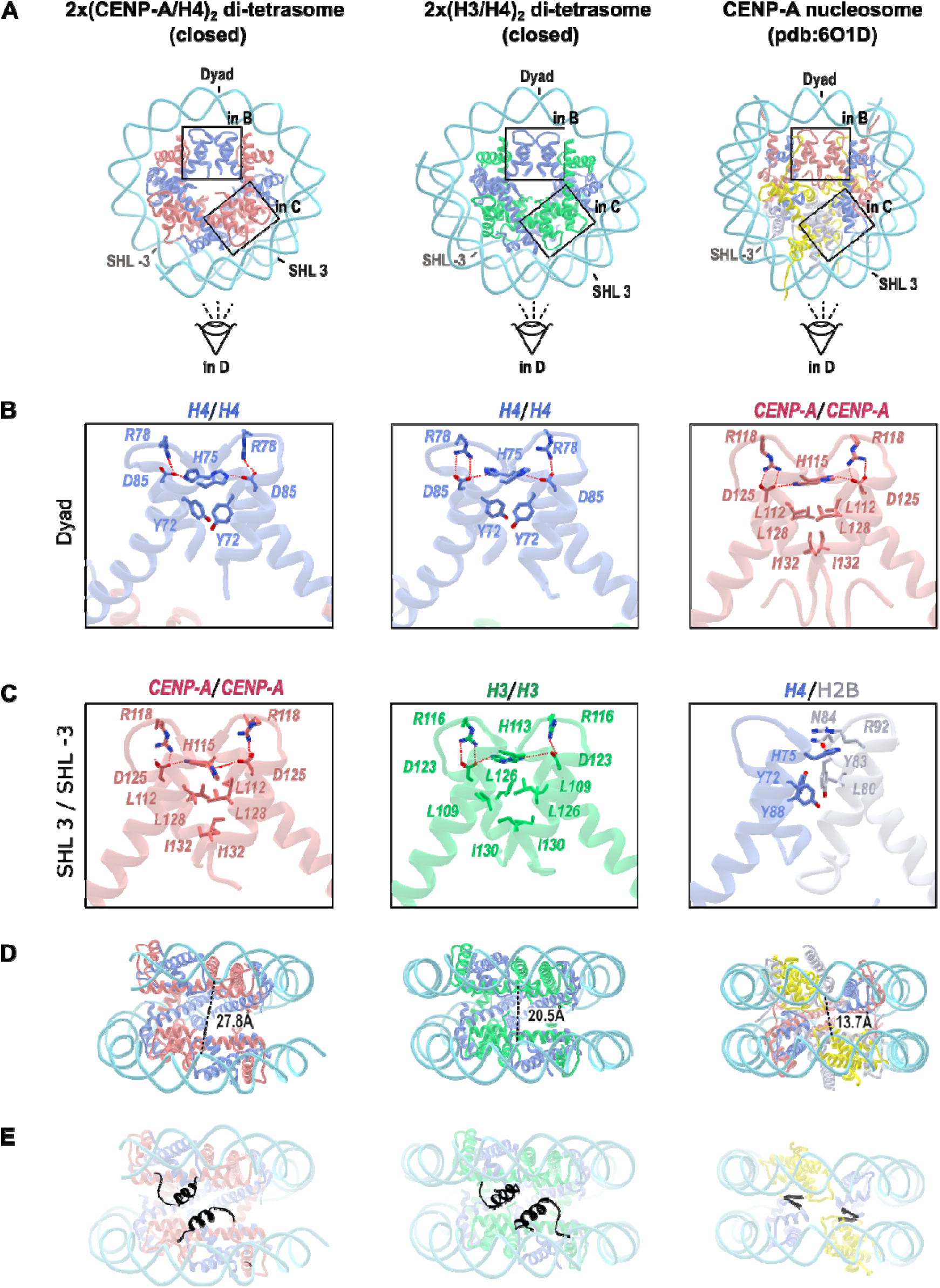
Di-tetrasomes have rotated H4/H4 four-helix bundle at the dyad and more separated DNA gyres. **A.** Ribbon representations of 2x(CENP-A/H4)_2_ di-tetrasome in closed conformation (left), 2x(H3/H4)_2_ di-tetrasome in closed conformation (middle) and CENP-A nucleosomes (PDB 6O1D) (right). CENP-A is red; H3 is green; H4 is blue; H2A is yellow; H2B is gray; and 145 bp DNA is cyan. The four-helix bundles at the dyad and at SHL3 are marked with rectangles and shown enlarged in B and C, respectively. The viewing direction for the side projection in panels D and E is indicated by an eye symbol. **B.** Enlarged view of the four-helix bundle at the dyad for the 2x(CENP-A/H4)_2_-di tetrasome (left), the 2x(H3/H4)_2_-di tetrasome (middle) and the CENP-A nucleosome (right). The key residues responsible for the interaction at the 4-helix bundles are shown in sticks. **C.** Enlarged view of the four-helix bundle at SHL3 and SHL-3 for the 2x(CENP-A/H4)_2_-di tetrasome (left), the 2x(H3/H4)_2_-di tetrasome (middle) and the CENP-A nucleosome (right). **D.** Side view of the particles from panel A (rotated clockwise by 90^0^ around the horizontal axis) to illustrate the separation of the DNA gyres. Measurements were made from base pairs labeled as −36 and 40 in chain J of DNA for di-tetrasomes and 36 and 114 on CENP-A nucleosome. **E.** Same as in D. Highlighted in black are the αN helices of CENP-A (left) and H3 (middle), which “push away” the two halves of the di-tetrasomes allowing for a more dynamic structure. On the right, the beta-sheet formed between H4 and H2A holds the CENP-A nucleosome (right) together to form a more compact particle.

Starting our analysis from the dyad, the di-tetrasomes accommodate H4/H4 four helix bundle at the dyad while nucleosomes have CENP-A/CENP-A or H3/H3 in this position. The hydrogen bonding network at the H3/H3 or CENP-A/CENP-A four helix bundle is robust and well preserved with strong hydrogen bonds ranging in size between 2.6 - 2.9 Å. The arginine and asparagine on one chain and histidine on the opposite chain are present and available for hydrogen bonding in both H3 and CENP-A interfaces and they form strong RDH-network: H3^Arg116^, H3^Asp123^ and H3^His113^ in canonical and CENP-A^Arg118^, CENP-A^Asp125^ and CENP-A^His115^ in CENP-A nucleosomes. Interestingly, similar RDH-network exists at the H4-H4 interface in di-tetrasomes (residues H4^Arg78^, H4^Asp85^ and H4^His75^) but here the distances between residues and their orientations are distorted (Figure 2B). In nucleosome and well as in di-tetrasome the rest of the interface at the dyad is mediated by hydrophobic residues (in nucleosome three leucine and one isoleucine while in di-tetrasome there are mainly aromatic residues (H4^Tyr72^, H4^His75^ and H4^Tyr88^). Because of loose hydrogen bonding, majority of interaction in H4/H4 interface at the dyad of di-tetrasomes are hydrophobic. This allows sliding at the four-helix bundle resulting in drastically different angles (∼40° rotation) between α2 helices (Figure S4A) in di-tetrasomes comparing to CENP-A nucleosomes.

Next, we analyzed the differences at the SHL3 and SHL-3 four-helix bundle interfaces (Figure 2C). Here, di-tetrasomes have CENP-A/CENP-A or H3/H3 four-helix bundles with weak RDH network, whereas nucleosomes have heterologous H4/H2B four-helix bundles mediated mainly by hydrophobic interactions and stacking of aromatic residues (H4^Y72^, H4^Y88^, H2B^Y83^ and H2B^L80^). Thus, the interface of the H4/H2B four-helix bundles exhibits fewer electrostatic interactions than the interfaces of H3/H3 or CENP-A/CENP-A at SHL3/SHL-3 in di-tetrasomes, suggesting that the compaction and stability of the nucleosome is not solely determined by interactions in the four-helix bundles. In fact, the distance between DNA gyres is much shorter in nucleosomes than in di-tetrasomes (Figure 2D). The increased compactness of the nucleosome is most likely caused by a beta-sheet within the histone core (highlighted in Figure 2E), which is formed between the beta-strand of H4 and the beta-strand of H2A. Here, the H4 residues (H4^96-98^) following the α3 helix of H4 form a beta-strand with the H2A residues (H2A^100-102^) following the α3 helix of H2A, while the remaining C-terminal residues of H2A are tightly stacked along the nucleosome face. The short beta-sheet between H4 and H2A holds the nucleosome together forming a stable particle. Interestingly, the same beta-sheet is reinforced (less solvent-exposed) in CENPA nucleosomes as an allosteric effect of CENP-C binding (56). This “stitching” interaction within the nucleosome is not present in di-tetrasomes. In addition, both H3- and CENP-A-containing di-tetrasomes lack the C-terminal tail of H2A, which could further stick the two halves of di-tetrasomes together. In contrast, both H3 and CENP-A have a long N-tail preceding to the α1 helix. This N-terminus contains a bulky, partially structured αN helix (not visible in our cryoEM data; highlighted in Figure 2E) facing the interior of a clam-shell-like structure that pushes away two halves of the particle.

The presence of bulky αN-helices of CENP-A that are opposing the dyad generates tension and leads to a rotation of the H4 four-helix bundle to a final 46° divergence with respect to the CENP-A four-helix bundles at the nucleosome dyad (Figure S4A). These changes contribute to a more “breathable” structure with increased flexibility and reduced DNA wrapping capacity, as reflected by more open and twisted DNA gyres on the opposite side of the dyad compared to a cannoncal nucleosome (Figure 2E). This is all consistent with the observation of open and closed conformations for di-tetrasomes, which are only two of many possible arrangements that we were able to refine from our cryo-EM data.

Thanks to this high flexibility, the di-tetrasomes can propagate on DNA to make longer “slinky-like” structures similarly as it was already reported for archaeal histones (57, 58). Indeed, by assembling (CENP-A/H4)_2_ on longer DNA, we have observed by cryoEM that (CENP-A/H4)_2_ tetramers can form “slinkies” (Figure S5; Table S2). Because di-tetrasomes are not so compact, the DNA at the entry/exit sites are further away from each other so DNA packed with di-tetrasome has distinguishingly different architecture then chromatin packed in nucleosomes (Figure S5D).

### 2x(CENP-A/H4)_2_ di-tetrasomes are stable but dynamic

To further investigate whether di-tetrasomes are stable, we performed molecular dynamics (MD) studies on CENP-A- and H3-containing di-tetrasomes (complete models with full-length histones and DNA) and nucleosomes (PDB 6O1D for CENP-A nucleosome and PDB 3LZ0 for H3 nucleosome) (Figure 3; Table S2). MD studies indicate that the di-tetrasomes are structurally stable but at the same time exhibit greater flexibility compared to nucleosome particles (Figure 3 and S6). The topological distribution of the most mobile regions in both the core histones and DNA is illustrated by the heat map, in which the residues that make up the protein/DNA complexes are colored according to their calculated B-factor (Figure 3A). The root mean square fluctuations (RMSF) for the histone fold domains (HFD) of di-tetrasomes are generally higher than those of nucleosomes. DNA generally fluctuates more than histones, both in di-tetrasomes and nucleosomes, and shows a tendency to unwrap from histone core at the DNA ends. In the HFD of 2x(CENP-A/H4)_2_ di-tetrasomes, (CENP-A/H4)_2_ dimers at the dyad (Figure S6A; chains C and E for CENP-A and D and F for H4) are more stable and comparable in fluctuations to dimers in 2x(H3/H4)_2_ di-tetrasomes, whereas (CENP-A/H4)_2_ dimers near the DNA ends (Figure S6A; chains A and G for CENP-A and B and H for H4) are very dynamic. Interestingly, RMSF analysis shows that the larger motions occur at the H4/H4 four-helix bundles at the dyad of di-tetrasomes (which includes L2 loops) and not at the CENP-A/CENP-A of H3/H3 four-helix bundles at the SHL-3 and SHL3 positions (which also include L2 of these histones). This is consistent with rotation at the dyad (Figure S4). In addition, great flexibility at the DNA level is also observed in di-tetrasomes compared to nucleosomes, with the largest amplitudes starting 50 bp away from the dyad. However, in nucleosomes, the rigidity of the HFD is transferred to the DNA regions that are in contact with the histones and show less variation than the parts exposed to the solvent. In contrast, the HFDs of di-tetrasomes are dominated by the fluctuations of the wrapped DNA. Overall, the fluctuations are higher in 2x(CENP-A/H4)_2_ di-tetrasomes compared to 2x(H3/H4)_2_ di-tetrasomes in both the protein and DNA parts.

**Figure 3.**
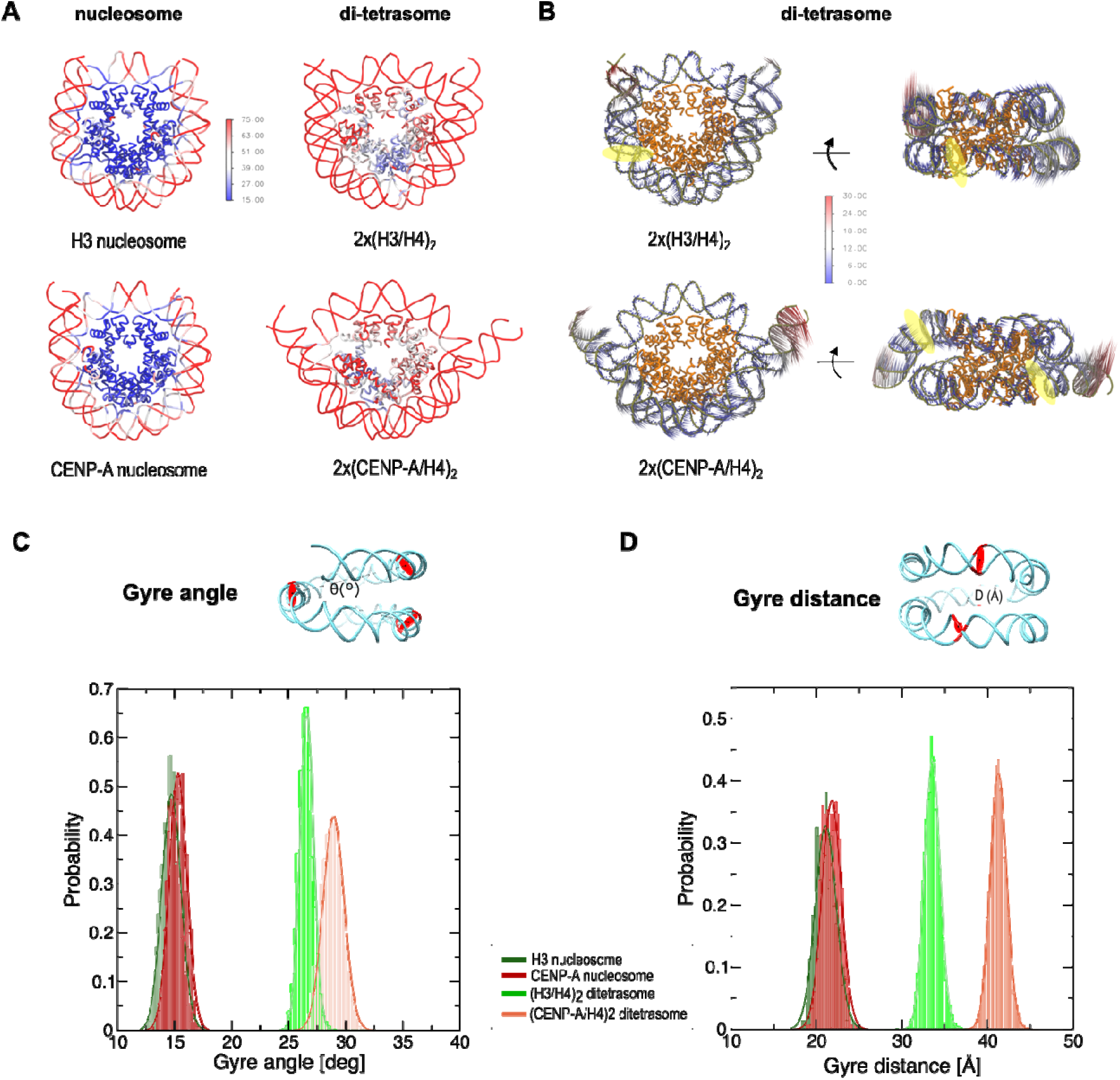
Molecular dynamics of 2x(CENP-A/H4)_2_ di-tetrasomes. **A.** The root-mean square fluctuations (RMSF) observed during MD simulations for nucleosome and di-tetrasomes is shown as a heat map on the ribbon structures of nucleosomes and di-tetraomes. The residues are colored according to their thermal B-factoras (blue – low fluctuations, red – high fluctuations) predicted by MD. **B.** The porcupine diagram illustrates the vectors of atomic displacements on the ribbon diagram of di-tetrasomes. The histone core is yellow, the DNA is gray. The arrows represent the directions and magnitudes of movement. The color code correlates with the size of the displacement in Å. **C and D.** Probability plots showing the distribution range of the opening angles (C) and the opening distances (D) of the DNA gyres for the di-tetrasome and nucleosomes structure observed in the MD simulation. The DNA base pairs (0, +39, −39) used to calculate the angles/distances are highlighted by red circles in the cyan ribbon diagram of DNA in 2x(CENP-A/H4)_2_ di-tetrasomes at the top of the plots.

To further analyze the concerted motions in di-tetrasomes (and nucleosomes), we performed principal component analysis (PCA). Indeed, the first eigenvector corresponds to an oscillatory movement of DNA ends around two hinge regions located at base pair positions (bps) −46 and +60 from the dyad in the 2x(CENP-A/H4)_2_-di tetrasome and at positions −39 and +60 from the dyad in the 2x(H3/H4)_2_-di tetrasome (yellow circles in Figure 3B). However, a similar movement was not observed in nucleosomes. The DNA deformation is accompanied by a corresponding movement in the L1-L2 region between chains C/D of the di-tetrasomes, which is localized near the DNA hinge. This is also very consistent with the experimental data which indicate multiple conformations after 52 bp from the dyad on one side and 62 bp on the other side, which could not be visualized in the cryoEM maps even after careful particle classification.

The deformation of DNA is not limited to the flexible ends, but also to the separation between the two gyres of the DNA double helix. Therefore, we measured clam-shell angle θ, which is defined between the center of mass of the DNA base pair at the dyad and the base pairs in the middle between the dyad and the DNA ends (Figure 3C). Here we see that both types of nucleosomes have θ = 18 ± 1° during the MD simulation, while 2x(H3/H4)_2_ di-tetrasomes have θ = 26 ± 1° and 2x(CENP-A/H4) di-tetrasomes have θ = 28 ± 1°. These values confirm that di-tetrasomes have a much larger gyre opening than nucleosomes and that the DNA gyres in 2x(CENP-A/H4)_2_ di-tetrasomes are significantly further apart compared to 2x(H3/H4)_2_ di-tetrasomes. A similar trend can be seen when measuring the distance between two gyres (Figure 3D; Table S3). It is noteworthy that both the distance between gyres and clam shell angle, although significantly different between the two di-tetrasomes types, are still quite stable and result in a relatively sharp peaks in the probability plot.

Overall, the MD analysis confirms that 2x(CENP-A/H4)_2_ di-tetrasomes are highly dynamic but stable molecular structures.

### (CENP-A/H4)_2_ tetrasomes and di-tetrasomes bind CENP-C with reduced affinity

Next, we wanted to understand if (CENP-A/H4)_2_ tetrasomes and 2x(CENP-A/H4)_2_ di-tetrasomes, as possible chromatin units, are able to recognize and bind CENP-A binding proteins, CENP-C and CENP-N.

It is known from previous structural work that two regions of human CENP-C (central binding domain, CENP-C^CR^ and CENP-C motif, CENP-C^motif^) bind CENP-A nucleosomes in a similar manner, with CENP-C^CR^ having higher affinity and specificity in the context of human proteins (8, 18, 59). When we incubate (CENP-A/H4)_2_ tetrasomes with CENP-C^CR^, we see clear binding (Figure 4A; the band corresponding to CENP-A tetrasomes disappears). In contrast, when (H3/H4)_2_ tetrasomes are incubated with CENP-C^CR^, CENP-C^CR^ binds to the free DNA to a higher extent than to H3 tetrasomes (Figure 4A). The band corresponding to free DNA disappears but the band corresponding to tetrasomes shows slightly decreased intensity with the addition of CENP-C^CR^. When 2x(CENP-A/H4)_2_ di-tetrasomes are incubated with CENP-C^CR^, we see clear binding for di-tetrasomes (the di-tetrasome band disappears), but unlike CENP-A nucleosome/CENP-C^CR^ complexes, which migrate as defined bands on the gel (Figure S7A), the binding of CENP-C^CR^ to di-tetrasomes results in a smear on the gel.

**Figure 4.**
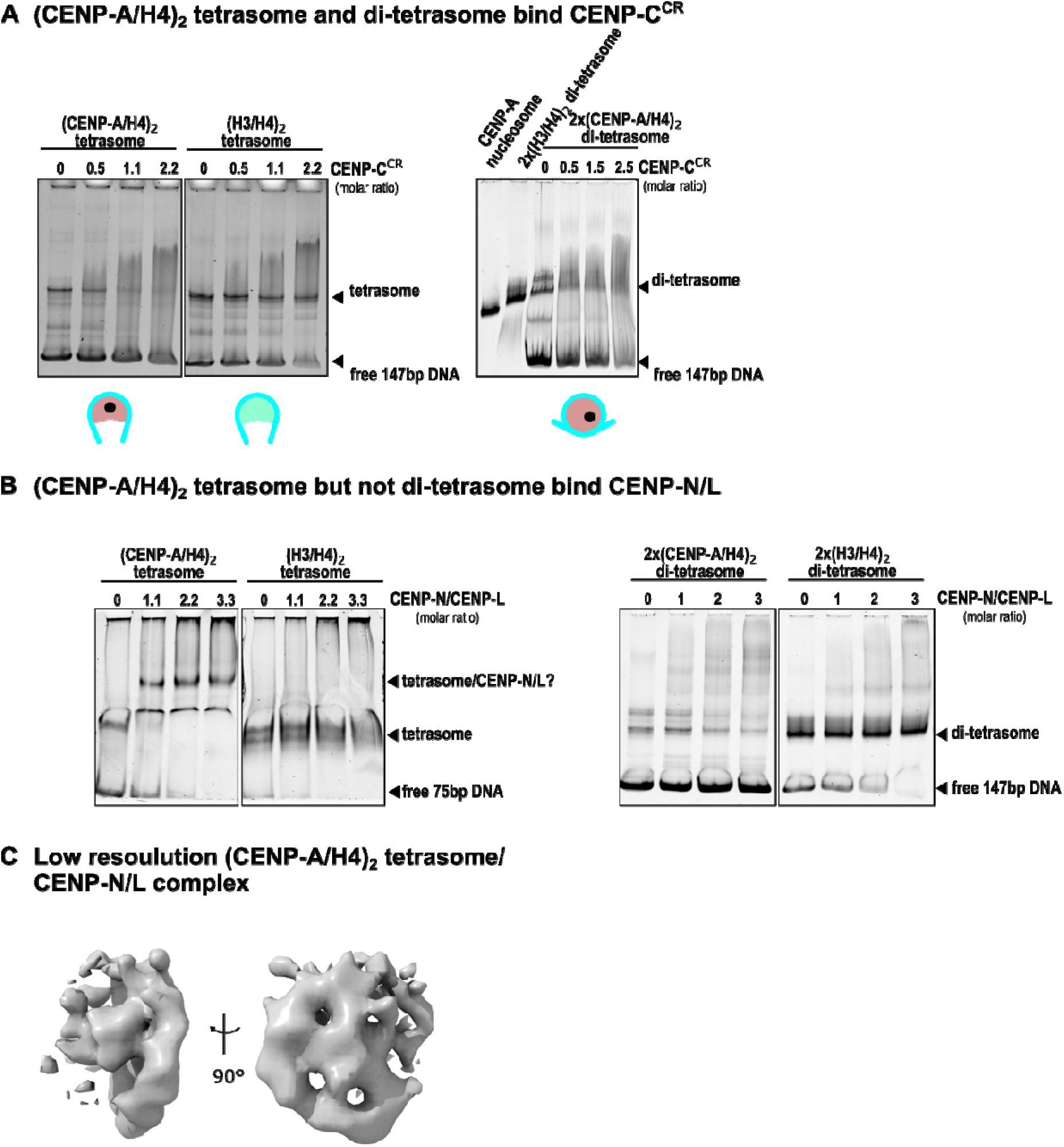
CENP-A-containing tetrasomes and di-tetrasomes bind CENP-C with low specificity but (CENP-A/H4)_2_ tetrasome binds CENP-N/L with high specificity. **A.** 5% native gels show that (CENP-A/H4)_2_ tetrasome (left) and di-tetrasome (right) bind CENP-C. Note that the band corresponding to the (di)-tetrasomes disappears, but the resulting complex runs as smear on the gel. This is in contrast to the binding of CENP-C by CENP-A nucleosomes, which results in complexes that run as single bands on the gel (see Figure S7A). Schematic representations of the tetrasomes and the 2x(CENP-A/H4)_2_ di-tetrasome are shown below the native gels, with the hydrophobic C-terminal tail of CENP-A shown as a black circle (on one side of the structure only). The CENP-A-containing histone core is red, the H3-containing histone core is green and the DNA is cyan. Note that the acidic patch of the H2A histone is missing in both tetrasomes and di-tetrasomes. **B.** 5% native gels show that the (CENP-A/H4)_2_ tetrasome binds CENP-N/L with high affinity, in contrast to the (H3/H4)_2_ tetrasome, which shows no binding (left gel). The gel on the right shows that the 2x(CENP-A/H4)_2_ di-tetrasome binds CENP-N/L only weakly, while the 2x(H3/H4)_2_ di-tetrasome does not bind it at all (right). **C.** Low-resolution cryo-EM map of the (CENP-A/H4)_2_ tetrasome:CENP-N/L complex shown on the native gel in (B).

The specific binding of CENP-C^CR^ by (CENP-A/H4)_2_ tetrasomes and di-tetrasomes but with lower affinity comparing to CENP-A nucleosomes is consistent with previous high-resolution structures of CENP-C^CR^/CENP-A nucleosome complexes (8, 11–13, 16, 19, 20). There, the binding is controlled by two interaction surfaces. One involves hydrophobic interactions between aromatic and hydrophobic residues in CENP-C^CR^ and the C-terminal part of CENP-A and the other interaction interface involves electrostatic interactions between two arginine residues in CENP-C and the acidic residues on H2A (Figure S7A). While both 2x(CENP-A/H4)_2_ di-tetrasomes and (CENP-A/H4)_2_ tetrasomes have a hydrophobic C-terminal part of CENP-A available for interaction with CENP-C^CR^, both lack the electrostatic part of the interactions provided by H2A. This would explain why CENP-A-containing complexes can still bind CENP-C^CR^, albeit with lower affinity compared to CENP-A nucleosomes.

### (CENP-A/H4)_2_ tetrasomes and di-tetrasomes bind N-terminal part of CENP-N

Although the binding of CENP-N to CENP-A was first demonstrated fifteen years ago (21), our understanding of this interaction has evolved with numerous high-resolution structures. Initial biophysical experiments and crystal structures involving only the CENP-A nucleosome and N-terminus of CENP-N, CENP-N^N-term^, indicated that CENP-N specifically recognizes the ^80^ArgGly^81^ insertion (CENP-A^RG-loop^) in histone CENP-A, which is absent in its canonical counterpart, H3. The CENP-A^RG-loop^ is exposed on the surface of CENP-A nucleosomes and is recognized and bound (together with the adjacent DNA) by several CENP-N residues (reviewed in (18)). However, full-length CENP-N forms a complex with CENP-L (60), and structures of CENP-A nucleosomes obtained in the presence of the full set of CCANs with fission yeast and human proteins, including the CENP-N/L complex, show that the CENP-N/L complex binds DNA extending from the nucleosome without specifically interacting with any histones in the nucleosomes. Moreover, the conformation that the CENP-N/L complex adopted in these structures is incompatible with the interaction involving the CENP-A^RG-loop^ in the context of the CENP-A nucleosome as observed in the CENP-A nucleosome/CENP-N^N-term^ structures (18). Different binding modes of CENP-N have raised the question of alternative arrangements between CENP-A and CCANs that could characterize different phases of the cell cycle (14). With this in mind, we tested whether (CENP-A/H4)_2_ tetrasomes and di-tetrasomes can bind CENP-N^N-term^ and the CENP-N/L complex.

First, we examined the binding of the N-terminal part of human CENP-N, CENP-N^N-term^, to (CENP-A/H4) tetrasomes and di-tetrasomes. We found that the band for 2x(CENP-A/H4)_2_ di-tetrasomes disappears from the gel even when substoichiometric amounts of CENP-N^N-term^ are added and several other bands are formed, which are likely to be complexes of CENP-N^N-term^ with DNA, tetrasomes and di-tetrasomes (Figure S7B, left gel). Interestingly, the binding of CENP-N^N-term^ to the CENP-A nucleosome, always leads to the formation of multiple laddered bands, indicating the formation of a CENP-A nucleosome/ CENP-N^N-term^ complex and its further oligomerization by CENP-N^N-term^ (Figure S7B, middle gel). This phenomenon was visualized before (24), but it is currently unclear whether the CENP-N-induced oligomerization has a physiological role in cells. Interestingly, we see that 2x(H3/H4)_2_ di-tetrasomes could also bind CENP-N^N-term^ and generate a band ladder on the native gel (Figure S7B, right gel).

### (CENP-A/H4)_2_ tetrasomes but not di-tetrasomes bind CENP-N/L complex

Since CENP-L, a binding partner of CENP-N, is also always present in centromeres, we tested whether (CENP-A/H4)_2_ tetrasomes and di-tetrasomes can bind the CENP-N/L complex (Figure 4B). Native gel analyzes show that 2x(CENP-A/H4)_2_ di-tetrasomes bind CENP-N/L in contrast to 2x(H3/H4)_2_ di-tetrasomes, but the binding is weak, and the resulting complexes are heterogeneous, leading to smearing on the gel. Interestingly, (CENP-A/H4)_2_ tetrasomes bind CENP-N/L very effectively and form a sharp, compact band on the gel. This is in contrast to (H3/H4)_2_-tetrasomes, which do not interact with the CENP-N/L complex. We also attempted to visualize the (CENP-A/H4)_2_/CENP-N/L complex by cryo-EM but could only obtain low-resolution density maps so far (Figure 4C).

Finally, we asked if (CENP-A/H4)_2_ tetrasomes could bind CENP-HIKM directly or upon binding CENP-N/L but the results were not conclusive.

## Discussion

Since the discovery that the position of the centromere on the chromosomes is dominated by an epigenetic rather than a genetic component (61), great efforts have been made to understand the molecular determinants of this phenomenon. In the following years, numerous experiments identified the histone H3 variant CENP-A as a key protein component that is necessary and sufficient for the establishment of functional centromeres (reviewed in (6)). However, only recently, thanks in part to rapid technological advances in cryo-EM, the high-resolution structure of CENP-A nucleosome in complex with CCANs from human and yeast has been published (11–13).

Surprisingly, however, there are no extensive contacts between protein part of the CENP-A nucleosome and the rest of the CCAN complex in the CENP-A/CCAN structure of *S. pombe* and humans. The only CCAN component that interacts with CENP-A is CENP-C, which remains unstructured, and thus “invisible”, in the cryoEM structure in the sections immediately before and after interaction with CENP-A (Figure 5 and S8). Furthermore, the same nucleosome/CCAN structure can also assemble without CENP-A, i.e. on naked DNA (10) or with the H3 nucleosome (13), so the question of how exactly CENP-A specifically recruits CCAN is therefore still open.

**Figure 5.**
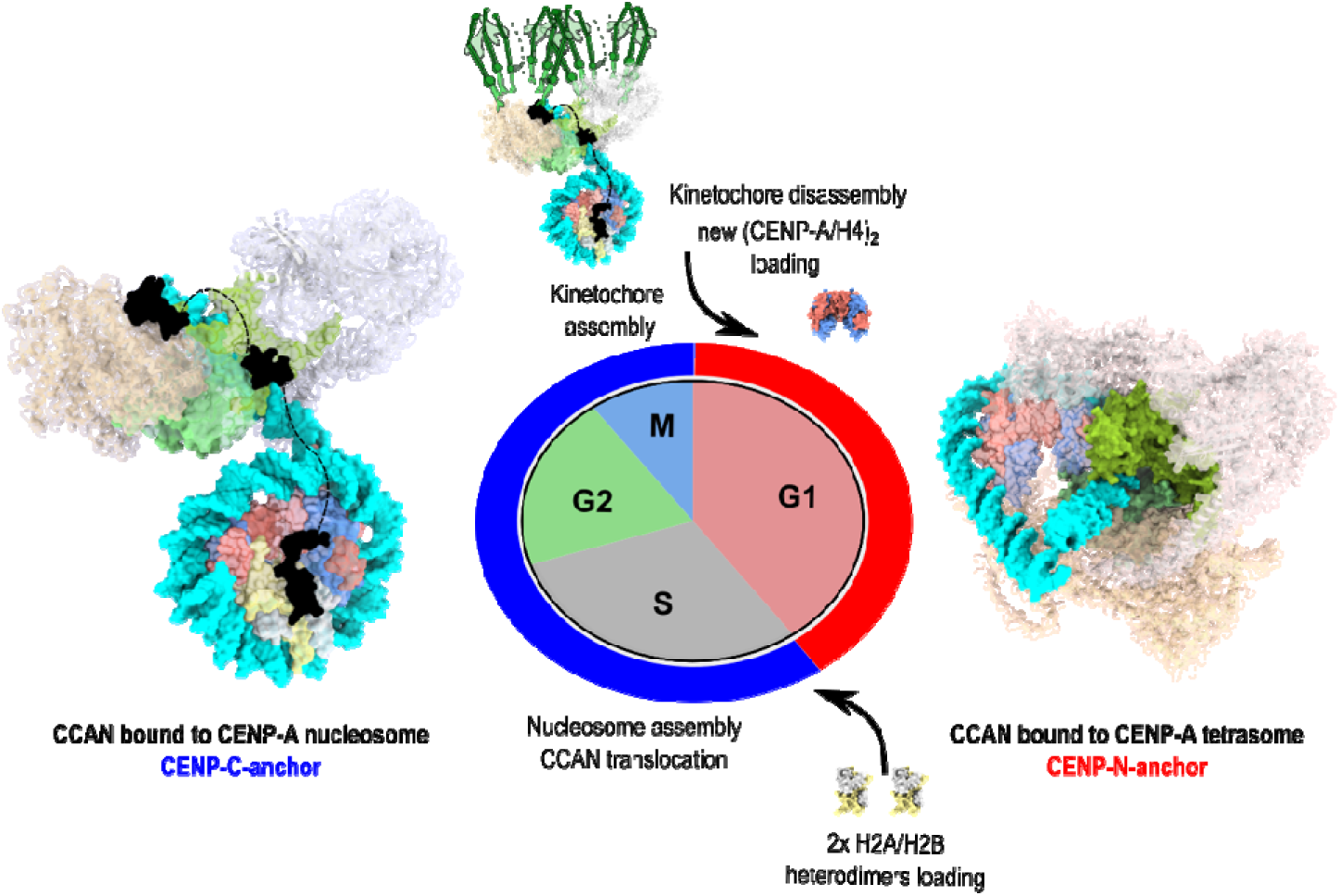
Model for cell cycle-dependent CENP-A: CCAN association alternating between CENP-C anchor and CENP-N anchor. In early G1 phase, after new CENP-A/H4 loading, the (CENP-A/H4)_2_ tetrasome structure is favored, and CCAN is bound to the CENP-A^RG loop^ with CENP-N as an anchor. In S phase, after H2A/H2B heterodimers are loaded onto the centromere, the CENP-A nucleosome is formed, and CCAN is translocated to the linker DNA and anchored to the CENP-A nucleosome by CENP-C. Space filling representation of CCAN bound to CENP-A nucleosome is generated using PDB 7YWX and CCAN bound to CENP-A tetrasome is generated using our structure of CENP-A tetrasome and AlphaFold modeling for CCAN complex. CENP-A is red, H4 is blue, H2A is yellow, H2B is grey, DNA is cyan, CENP-C is black, CENP-N-L is green, CENP-H-I-K-M is beige, and CENP-O-P-Q-U-R is light gray.

Here we present the structure of 2x(CENP-A/H4)_2_ di-tetrasomes that could form in chromatin in the absence of H2A/H2B dimers. The structures are similar to already reported 2x(H3/H4)_2_ di-tetrasomes or “H3-H4 octasome”, whose cryoEM structure and possible existence within the cell has been demonstrated by cross-linking experiments (55). More recently, a creative experiment demonstrated that the *E. coli* genome can be “packaged” in functional *E.coli* cells expressing only histones H3 and H4 (62) confirming that (H3/H4)_2_ tetrasomes and di-tetrasomes can be formed in living cells. Also noteworthy is the observation of more flexible chromatin particles with a wider opening of DNA gyres visualized by cryotomography on chromatin from interphase nuclei (63), suggesting more dynamic nucleosomes in the cell context and also the possible presence of di-tetrasomes. In addition, we have also observed that the 2x(CENP-A/H4)_2_ di-tetrasomes when tightly packed, form a peculiar shape of nucleosome fibers similar to that reported for archaeal histones (57, 58), and this “looser” flexible packing may also contribute to a specific chromatin feature characteristic of centromeres.

Our data suggest that (CENP-A/H4)_2_ tetrasomes bind CENP-N/L strongly, probably involving the CENP-A^RG loop^, and bind CENP-C poorly because they lack the H2A acid patch. In contrast, CENP-A nucleosomes bind CENP-C strongly but cannot bind CENP-N/L specifically via the CENP-A^RG-loop^ because CCAN complex sterically collide with the second DNA wrap harboring H2A/H2B. This implies that (CENP-A/H4)_2_ (di)-tetrasomes could serve as the basis for kinetochore organization in the absence of H2A/H2B and through CENP-A^RG-loop^ CENP-N/L interactions.

We propose that CENP-A alternates between (CENP-A/H4)_2_ (di)-tetrasomes and CENP-A nucleosomes during the cell cycle with two different CCAN binding modes, one involving the CENP-A^RG-loop^ and using CENP-N as an anchor and the other using CENP-A^C-term^ and H2A^acidic patch^ and CENP-C as an anchor (Figure 5). Plastic kinetochores that switch between CENP-A/CENP-C anchor in mitosis and CENP-A/CENP-N anchor in interphase were previously proposed by the Fukagawa lab based on the observation that binding of phosphorylated chicken CENP-C occurs in a conformation that hinders interaction with the CENP-A^RG-loop^ (19, 64). Here, we extend this proposal suggesting that CENP-A exists in the form of (CENP-A/H4)_2_ tetrasomes and di-tetrasomes during the loading of new CENP-A in G1 phase of the cell cycle and matures into CENP-A nucleosome during S-phase when bulk of H2A/H2B dimers are being incorporated in chromatin.

CENP-A loading is known to occur upon exit from mitosis (65) and not during DNA replication, as is the case for bulk histones, and two very recent studies have revealed molecular details of the spatiotemporal mechanisms behind the deposition of new CENP-A (66, 67). Last step in the process of CENP-A deposition is done by histone chaperone HJURP (68, 69) that deposits CENP-A/H4 dimer into centromeric chromatin (70–73), but the exact mechanism of how this histone pair is incorporated into DNA is still unclear.

It has been proposed that H3.3 is loaded into centromeric chromatin in S phase as a “placeholder” and then replaced by CENP-A at the exit of mitosis (74), but it is not clear how H3.3 is directed to specific sites in centromeric chromatin and whether the replacement in G1 involves the entire nucleosome or only the subnucleosomal complexes. Interestingly, Bodor et al. (75) found in a follow-up study that CENP-A is loaded in early G1 together with H4, but without concomitant loading of H2A/H2B. Moreover, the CENP-A/H4 subnucleosomal core remains stably incorporated over multiple cell divisions, whereas all other histones turn over quite rapidly. The exceptional chromatin stability of the CENP-A/H4 subnucleosomal core is directed by the CENP-A Targeting Domain (CATD) (76). The CATD comprises the L1 loop (containing CENP-A^RG-loop^) and the α2 helix of CENP-A and is required for the binding of HJURP and targeting of CENP-A to the centromere (76, 77). However, the chromatin stability of CENP-A/H4 observed by Bodor et al. (75) was not found to be dependent on HJURP. This result could be explained in part with strong hydrophobic association between H4 and CENP-A (25, 77) and with our model where, immediately after chromatin incorporation, (CENP-A/H4)_2_ tetrasomes bind the CCAN via CENP-N and through the CENP-A^RG-loop^. Indeed, we created an AlphaFold model that has (CENP-A/H4)_2_ tetrasome as a base and harbors CCAN through CENP-A^RG-loop^ interaction (Figure 5 and Figure S8B), and we see no steric clashes. Furthermore, in the model, the CCAN tightly wraps around (CENP-A/H4)_2_ tetrasome, shielding it from other possible interactions. This interaction would ensure the stability of CENP-A/H4 and the CCAN at the centromere before the S phase. During the S phase, CENP-A nucleosome is completed by the incorporation H2A/H2B, which allows the translocation of the CCAN to nucleosomal DNA (potentially by remodelers (78)), where it stays anchored by CENP-C and ready to recruit kinetochore in mitosis.

Although theoretically possible, we were unable to successfully assemble the (CENP-A/H4)_2_-tetrasome-CCAN complex *in vitro* with recombinantly purified CCAN, suggesting that such a structure, if indeed present in the cell, would need to be further stabilized by components missing from our *in vitro* assembly or by possible posttranslational modifications of the CCAN components and/or CENP-A. We also found that 2x(CENP-A/H4)_2_ di-tetrasomes readily convert to more stable nucleosomes in the presence of free H2A/H2B, which would complicate the isolation of such particles from cells and would be consistent with a previous detailed study in which full octameric homotypic CENP-A nucleosomes were found present throughout the cell cycle upon chromatin isolation (27).

Although we are aware that our model is speculative in the absence of direct cell-based evidences, we believe that our results will motivate further research to answer two important questions: 1. how are the centromere proteins organized immediately after CENP-A loading in G1 phase? and 2. when and how (or if at all) does the specific interaction between the CENP-A^RG-loop^ and CENP-N/L take place during the cell cycle? We believe that the further development of imaging techniques such as cryo-electron tomography, which allow visualization of centromeres in natural environment (79), and the isolation of native kinetochores (80), are crucial to obtain an accurate and detailed picture of centromere organization during the cell cycle.

## Data availability

Atomic coordinate model for 2x(CENP-A/H4)_2_ di-tetrasome have been deposited in the PDB with accession code 9GXA. CryoEM maps have been deposited in the EMBD with the accession codes: EMD-51645, EMD-51646, EMD-51647, EMD-51656.

Molecular dyanamics data for di-tetrasomes is available at https://archive.sigma2.no/pages/public/datasetDetail.jsf?id=10.11582/2024.00125 (doi: 10.11582/2024.00125) and for nucleosomes at https://archive.sigma2.no/pages/public/datasetDetail.jsf?id=10.11582/2024.00135 (doi: 10.11582/2024.00135).

## Acknowledgments and funding

We thank Prof. Lars Jansen (University of Oxford) for discussions at the beginning of the project phase and all members of the Sekulic group for their support and discussions throughout the project. NS and AAA are supported by the NCMM and the Research Council of Norway (grant numbers 187615 and 325528). M.M., X.L., Ma.C., Mi.C. were supported by the Research Council of Norway through the Centre of Excellence Hylleraas Centre for Quantum Molecular Sciences (grant number 262695), and the Norwegian Supercomputing Program (NOTUR) for computing hours and storage facilities (grants numbers NN4654K and NS4654K). Work in the Halic laboratory is funded by St. Jude Children’s Research Hospital, the American Lebanese Syrian Associated Charities, and NIH awards 1R01GM135599 and 1R01GM141694. For the initial protein expressions in insect cells, we used the protein purification facility LP3 at Lund University, which is funded by the Swedish Research Council as a national research infrastructure. The cryo-EM data were collected at the Umeå Core Facilities for Electron Microscopy, a node of the Cryo-EM Swedish National Facility, which is funded by the foundations of Knut and Alice Wallenberg, Erling Persson and the Kempe family, SciLifeLab, Stockholm University and Umeå University.

## Author contributions

NS and AAA conceived the project. AAA purified proteins and DNA and assembled the complexes. AAA performed all binding assays and MNase assays. AAA prepared the samples for CryoEM and processed the CryoEM data. MH and SB helped with cryoEM data processing and interpretation. MM performed all MD experiments under the supervision and guidance of XL, Ma.C., Mi.C. NS and AAA analyzed and interpreted the CryoEM data and wrote the manuscript with the input from all authors.

