## Supplementary material for "Non-nucleosomal (CENP-A/H4)_2_ - DNA complexes as a possible platform for centromere organization" (from another version of this preprint: version 1, 2025-01-01)

<u>This supplementary file contains:</u>

Supplementary tables S1 to S4

Supplementary figures S1 to S8

**Table S1. Cryo-EM data collection statistics for 2x(CENP-A/H4)$_2$ and 2x(H3/H4)$_2$ di-tetrasomes**

| | | |
|---|---|---|
| **Data collection** | | |
| Magnification | 165 000x | |
| Voltage (kV) | 300 | |
| Electron exposure (e–/Å$^2$) | 57.5 | |
| Defocus range in µm (steps) | -1.5 – -3.0 (0.3) | |
| Pixel size (Å) | 0.82 | 0.82 |
| **2x(H3/H4)$_2$ $^{\alpha\text{-sat}}$ di-tetrasome data processing** | **Open conformation**<br>EMD-51647<br>(Most unwrapped) | **Closed conformation**<br>EMD-51646<br>(Most wrapped) |
| Initial reference map | ab initio | ab initio |
| Symmetry imposed | C1 | C1 |
| Initial particle images (no.) | 158.3 K | 158.3 K |
| Final particle images (no.) | 64.7 K | 57.5 K |
| Box size (Å) | 360 | 360 |
| Pixel size (Å) | 0.82 | 0.82 |
| Map resolution (Å), FSC threshold | 3.75 | 3.82 |
| **2x(CENP-A/H4)$_2$$^{\alpha\text{-sat}}$ di-tetrasome data processing** | **Closed conformation**<br>EMD-51645<br>PDB: 9GXA<br>(Most wrapped) | |
| Initial reference map | ab initio | |
| Symmetry imposed | C1 | |
| Initial particle images (no.) | 119.3 K | |
| Final particle images (no.) | 22.5 K | |
| Box size (Å) | 360 | |
| Pixel size (Å) | 0.82 | |
| Map sharpening *B* factor (Å$^2$) | -84 | |
| Map resolution (Å), FSC threshold | 4.01 | |
| **Refinement** | | |
| Initial model used | - | |
| Model resolution (Å) FSC threshold | 4 | |

| | |
|---|---|
| Model composition | |
| Nonhydrogen atoms | 9226 |
| Protein residues | 575 |
| Nucleotides | 224 |
| Ligands | 0 |
| R.m.s. deviations | |
| Bond lengths (Å) | 0.004 |
| Bond angles (°) | 0.652 |
| Validation | |
| MolProbity score. | 1.7 |
| Clashscore | 15.93 |
| Poor rotamers (%) | 0 |
| Ramachandran plot | |
| Favored (%) | 98.39 |
| Allowed (%) | 1.61 |
| Disallowed (%) | 0 |

**Table S2. Cryo-EM data collection statistics for (CENP-A/H4)$_2$ poly-tetrasome.**

| **Data collection** | |
|---|---|
| Magnification | 165 000x |
| Voltage (kV) | 300 |
| Electron exposure (e–/Å$^2$) | 50 |
| Defocus range in µm (steps) | -0.8 – -2.4 (0.3) |
| Pixel size (Å) | 0.704 |
| **4x(CENP-A/H4)$_2$ $^{di\text{-}\alpha\text{-}sat}$ tetra-tetrasome data processing** | EMD-51656 |
| Initial reference map | ab initio |
| Symmetry imposed | C1 |
| Initial particle images (no.) | 76.8 K |
| Final particle images (no.) | 27 K |
| Box size (Å) | 280 |
| Pixel size (Å) | 1.3074 |
| Map resolution (Å), FSC threshold | 7.01 |

**Table S3. Number of atoms composing the simulated systems, and total simulation time.**

| | Protein/DNA | Water | $Na^+$ | $Cl^-$ | Total | Simulated time |
|---|---|---|---|---|---|---|
| 3LZ0 | 24’751 | 757’812 | 864 | 723 | 784’150 | 900 ns |
| 6O1D | 25’862 | 756’285 | 865 | 724 | 783’736 | 900 ns |
| $2x(H3/H4)_2$ | 24’936 | 757’368 | 860 | 719 | 783’883 | 900 ns |
| $2x(CENP\text{-}A/H4)_2$ | 25’236 | 756’900 | 872 | 731 | 783’739 | 900 ns |

**Table S4. Gyre data.**

| | Gyre distance<br>Grye angle<br>(MD data) | Gyre distance<br>Gyre angle<br>(PDB) |
|---|---|---|
| 3LZ0 | 21.1 ± 1.2 Å<br>14.7 ± 0.8 deg | 22.7 Å<br>15.7 deg |
| 6O1D | 21.8 ± 1.1 Å<br>15.3 ± 0.8 deg | 26.4 Å<br>17.9 deg |
| $2x(H3/H4)_2$ | 33.5 ± 0.8 Å<br>26.6 ± 0.6 deg | 36.8 Å<br>26.4 deg |
| $2x(CENP\text{-}A/H4)_2$ | 41.3 ± 1.0 Å<br>28.9 ± 0.9 deg | 40.2 Å<br>28.4 deg |

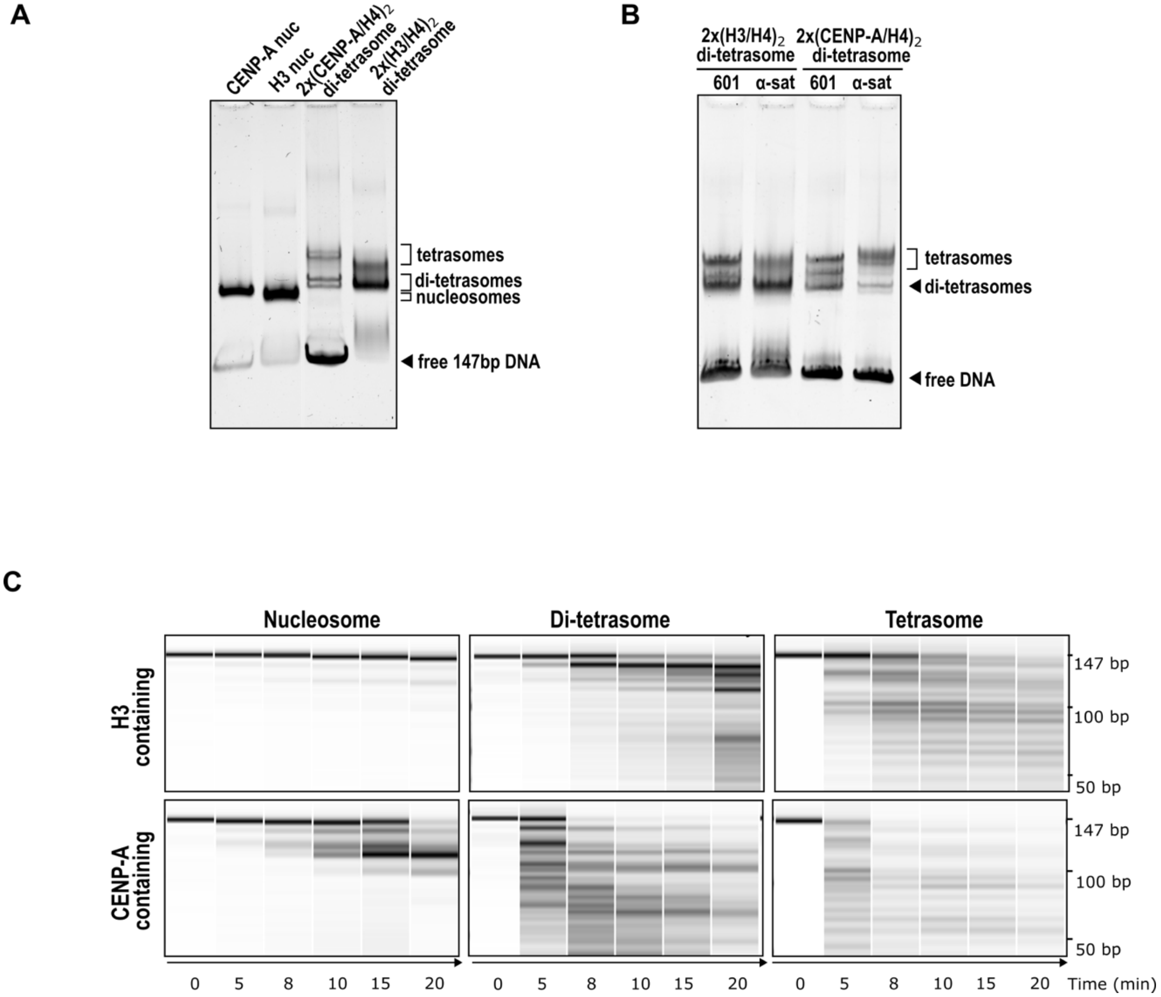


**Figure S1. 2x(CENP-A/H4)$_2$ tetramers wrap the DNA and form flexible di-tetrasome. A.** 5% native gel showing migration of assembled H3 and CENP-A nucleosomes and 2x(H3/H4)$_2$ and 2x(CENP-A/H4)$_2$ di-tetrasomes. We also observe that (H3/H4)$_2$ tetramers stay bound to DNA in tetrasome or di-tetrasome form while running on the gel, while (CENP-A/H4)$_2$ are generally less stable, resulting in large amounts of unassembled DNA. **B.** 2x(H3/H4)$_2$ and 2x(CENP-A/H4)$_2$ tetrasomes and di-tetrasomes form on both super positioning 601 and alpha-satellite DNA. **C.** Virtual gels from Bioanalyzer showing Micrococcal nuclease digestion of DNA when assembled with H3 (upper gels) or CENP-A (lower gels) and appropriate histones to form nucleosome (left), di-tetrasome (middle) and tetrasome (right).

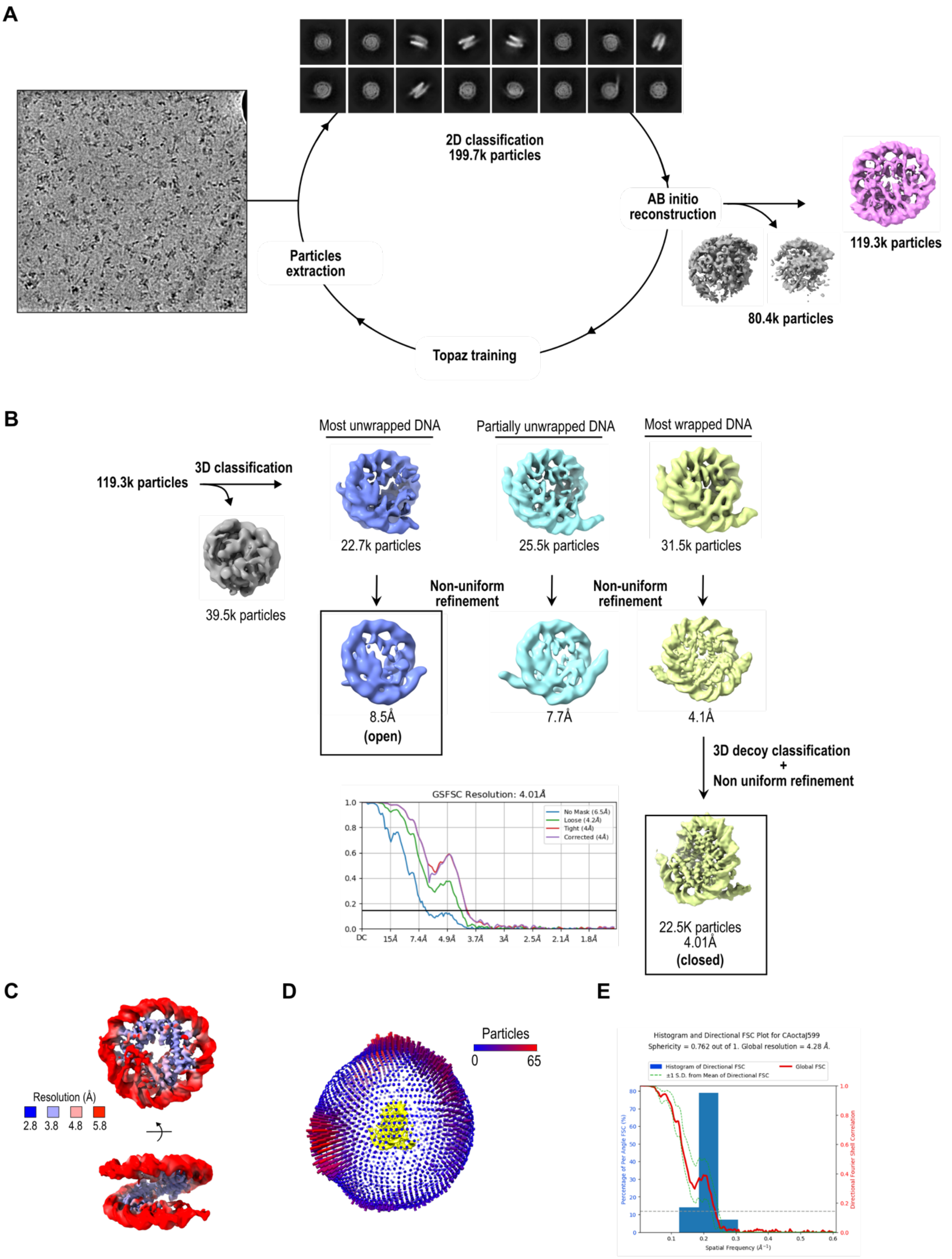
A
2D classification
199.7k particles
AB initio reconstruction
119.3k particles
80.4k particles
Particles extraction
Topaz training
B
Most unwrapped DNA
Partially unwrapped DNA
Most wrapped DNA
3D classification
119.3k particles
39.5k particles
22.7k particles
25.5k particles
31.5k particles
Non-uniform refinement
Non-uniform refinement
8.5Å
(open)
7.7Å
4.1Å
3D decoy classification
+
Non uniform refinement
GSFSC Resolution: 4.01Å
No Mask (6.5Å)
Loose (4.2Å)
Tight (4Å)
Corrected (4Å)
22.5K particles
4.01Å
(closed)
C
Resolution (Å)
2.8 3.8 4.8 5.8
D
Particles
0 65
E
Histogram and Directional FSC Plot for CAoctaJ599
Sphericity = 0.762 out of 1. Global resolution = 4.28 Å.
Histogram of Directional FSC
±1 S.D. from Mean of Directional FSC
Global FSC
Percentage of Per Angle FSC (%)
Directional Fourier Shell Correlation
Spatial Frequency (Å⁻¹)

**Figure S2. Cryo-EM analysis of 2x(CENP-A/H4)$_2$ di-tetrasome. A.** From left to right: Representative cryo-EM micrograph, subset of selected 2D class averages and the reference map for 2x(CENP-A/H4)$_2$ di-tetrasome generated using *ab initio* reconstruction. Discarded classes from ab initio reconstruction are shown in grey. **B.** Flowchart of 3D classification analysis for 2x(CENP-A/H4)$_2$ di-tetrasome. After homogeneous refinement of the selected particles from *ab initio* reconstruction, 3D classification followed by heterogeneous refinement was performed to classify the differences at the DNA entry/exit site. Classes are grouped according to DNA conformation at the entry/exit sites. Heterogeneous refinement is followed by non-uniform refinement, and the particles from the class with highest resolution were further processed using 3D decoy classification and non-uniform refinement. The FSC cruve is shown next to the final map. Discarded classes are shown in grey. **C.** Local resolution map of CENP-A di-tetrasome. **D.** Euler angle distribution of particles used in the 3D reconstruction of the most wrapped class. **E.** Histogram of directional FSC plot showing uniform resolution in all directions and a sphericity of 0.762.

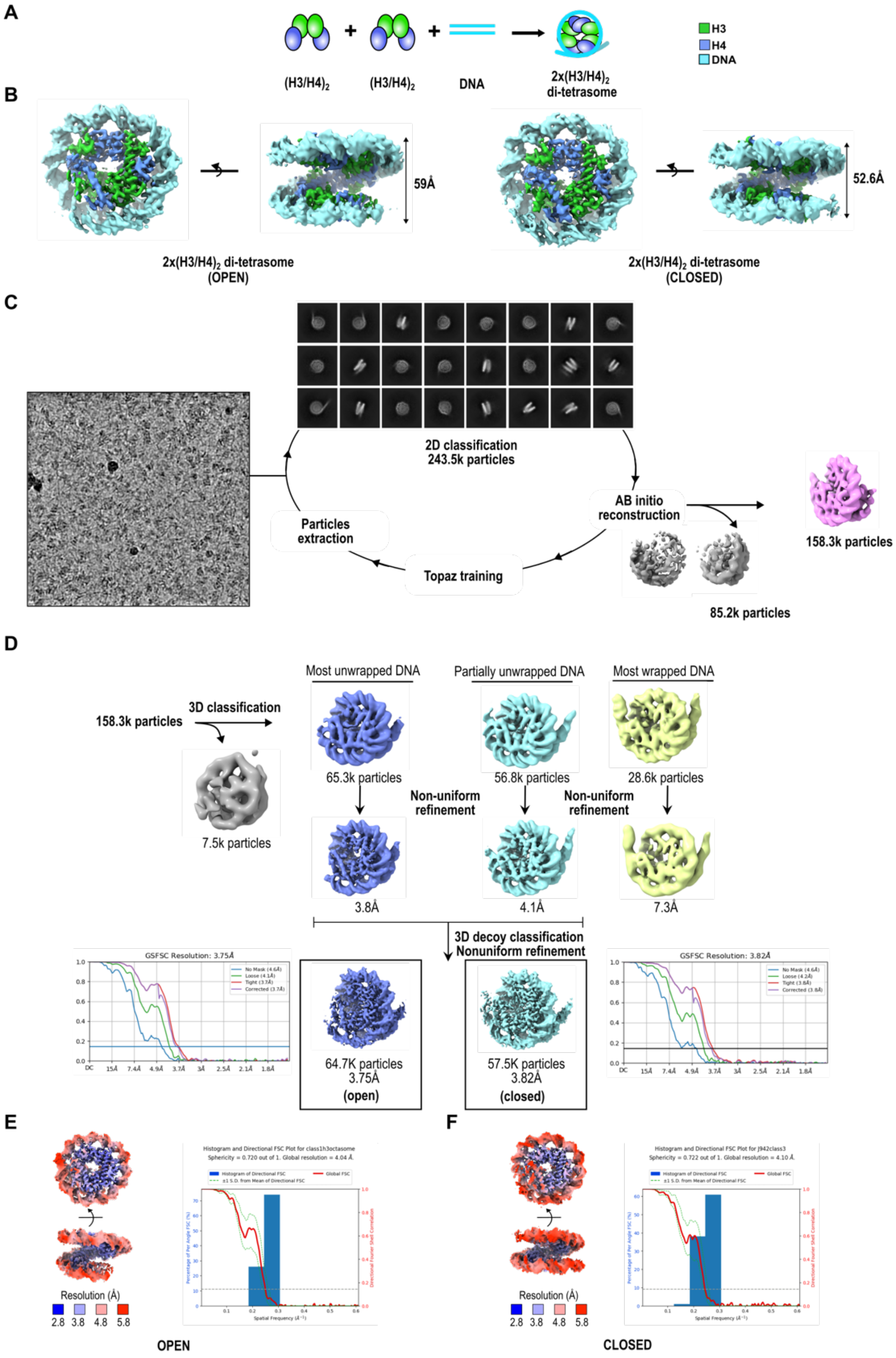
A
H3
H4
DNA
(H3/H4)2
(H3/H4)2
DNA
2x(H3/H4)2 di-tetrasome
B
59Å
52.6Å
2x(H3/H4)2 di-tetrasome (OPEN)
2x(H3/H4)2 di-tetrasome (CLOSED)
C
2D classification 243.5k particles
AB initio reconstruction
158.3k particles
Particles extraction
Topaz training
85.2k particles
D
158.3k particles
3D classification
7.5k particles
Most unwrapped DNA
Partially unwrapped DNA
Most wrapped DNA
65.3k particles
56.8k particles
28.6k particles
Non-uniform refinement
Non-uniform refinement
3.8Å
4.1Å
7.3Å
3D decoy classification
Nonuniform refinement
GSFSC Resolution: 3.75Å
64.7K particles
3.75Å
(open)
57.5K particles
3.82Å
(closed)
GSFSC Resolution: 3.82Å
E
F
Resolution (Å)
2.8 3.8 4.8 5.8
OPEN
Resolution (Å)
2.8 3.8 4.8 5.8
CLOSED

**Figure S3. Cryo-EM analysis of 2x(H3/H4)$_2$ di-tetrasome. A.** Schematic representation of assembly of 2x(H3/H4)$_2$ di-tetrasomes. H3 is green; H4 is blue; H2A is yellow; H2B is gray; and 145 bp DNA is cyan. **B.** cryoEM maps for H3 di-tetrasome in open (left) and closed (right) conformations with measurements of the opening of the DNA gyres on the opposite side of the dyad. Histones and DNA are colored as in A. **C.** From left to right: Representative cryo-EM micrograph, subset of selected 2D class averages and the reference map for 2x(H3/H4)$_2$ di-tetrasome generated using *ab initio* reconstruction. Discarded classes from *ab initio* reconstruction are shown in gray. **D.** Flowchart of 3D classification analysis for 2x(H3/H4)$_2$ di-tetrasome. After homogeneous refinement of the selected particles from *ab initio* reconstruction, 3D classification followed by heterogeneous refinement was performed to classify the differences at the DNA entry/exit site. Classes are grouped according to DNA conformation at the entry/exit sites. Heterogeneous refinement is followed by non-uniform refinement, and the particles from the class with highest resolution were further processed using 3D decoy classification and local refinement. The FSC curves from the non-uniform refinement are shown next to the final maps. Discarded classes are shown in gray. **E-F.** Local resolution map and the histogram of directional FSC plot showing uniform resolution in all directions for the open (E) and closed (F) conformations of (H3/H4)$_2$ di-tetrasome with a sphericity of 0.72.

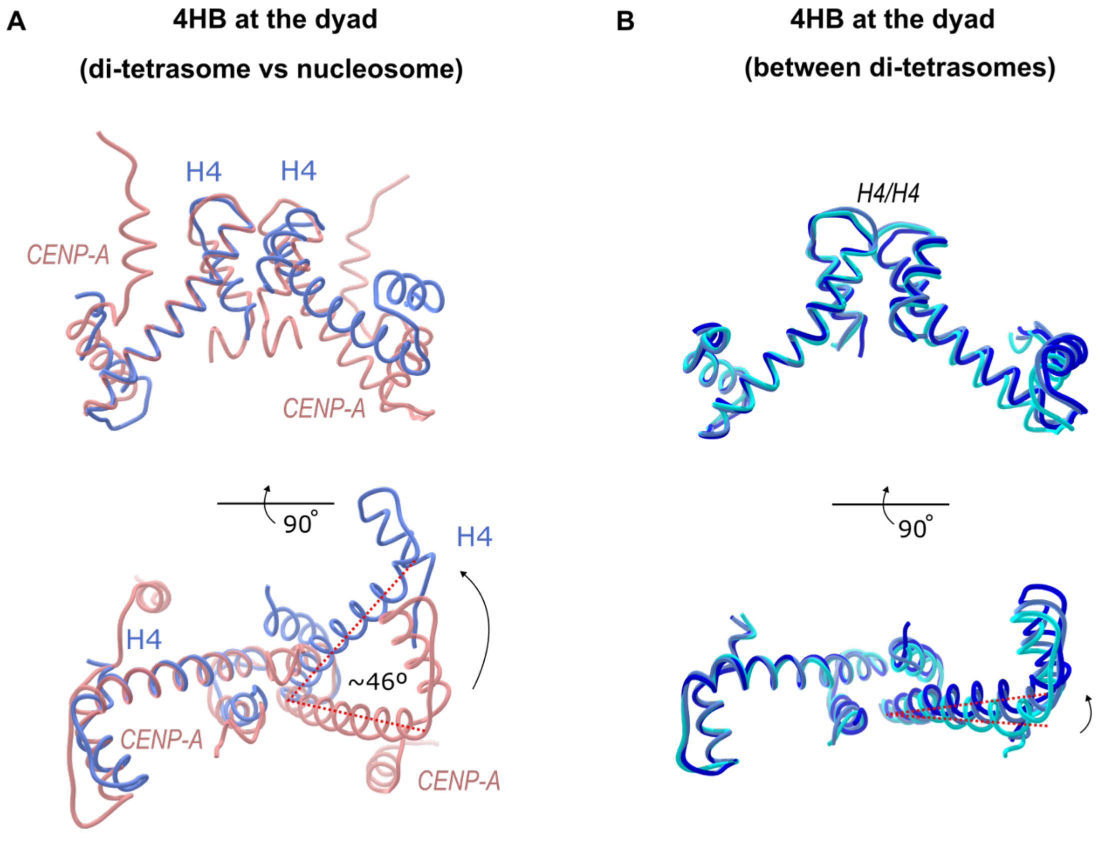


| | angle (°) |
|---|---|
| CENP-A nucleosome (6O1D) | 0 |
| 2x(CENP-A/H4)$_2$ di-tetrasome (closed) | 46 |

| | angle (°) |
|---|---|
| 2x(H3/H4)$_2$ di-tetrasome (closed) | 0 |
| 2x(H3/H4)$_2$ di-tetrasome (open) | 4 |
| 2x(CENP-A/H4)2 di-tetrasome (closed) | 6 |

**Figure S4. Di-tetrasomes have rotated H4/H4 four-helix bundle at the dyad. A.** Overlay of four-helix bundle (4HB) at the dyad of CENP-A di-tetrasome (H4/H4 interface; blue) in comparison to CENP-A nucleosome (PDB 6SE0; CENP-A/CENP-A interface; red) showing ~46$^0$ degrees rotation. **B.** Overlay of 4HB at the dyad of 2x(CENP-A/H4)$_2$ di-tetrasome in comparison to the two classes (open and closed) of 2x(H3/H4)$_2$ di-tetrasome showing flexible interface.

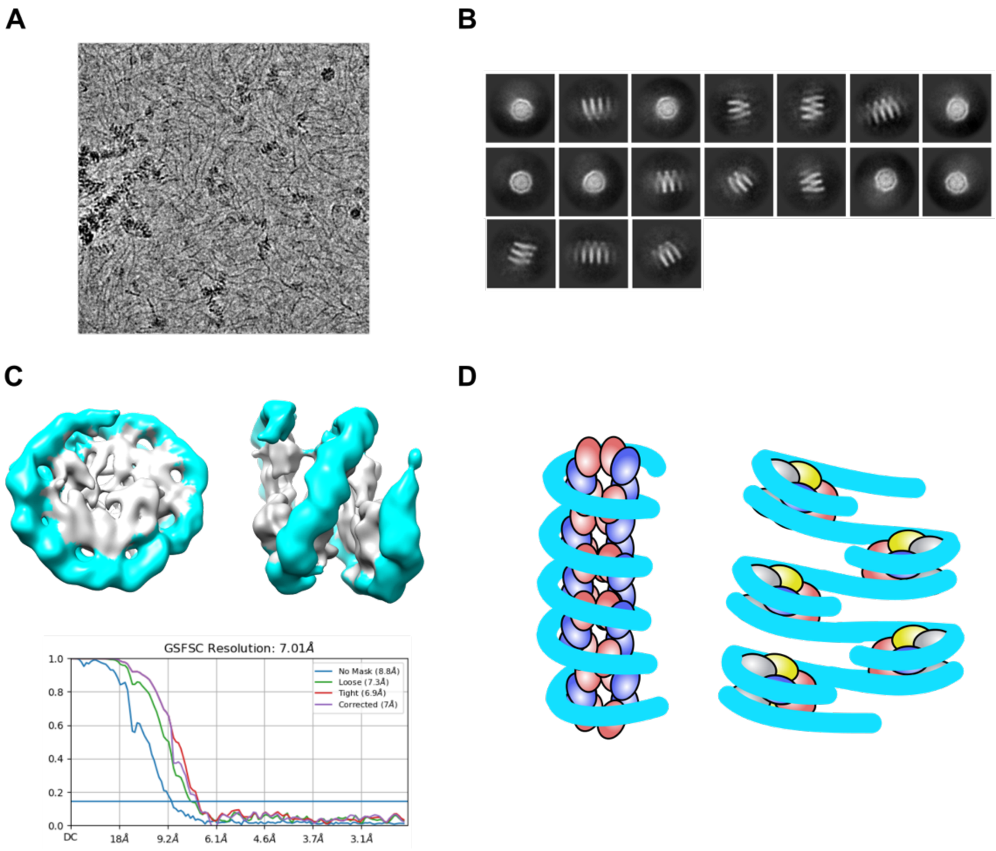


**Figure S5. (CENP-A/H4)$_2$ di-tetrasomes "stack" to form a unique chromatin architecture. A.** Representative cryo-EM micrograph of CENP-A tetra-tetrasome assembled on 342 bp di-$\alpha$-satellite DNA. **B.** Subset of selected 2D class averages. **C.** Cryo-EM map of the tetra-tetrasome reconstructed using non-uniform refinement and the corresponding FSC curve. **D.** A model of tetrasome-loaded chromatin versus nucleosome-loaded chromatin CENP-A is red, H4 is blue, H2A in yellow, H2B in grey and DNA is cyan.

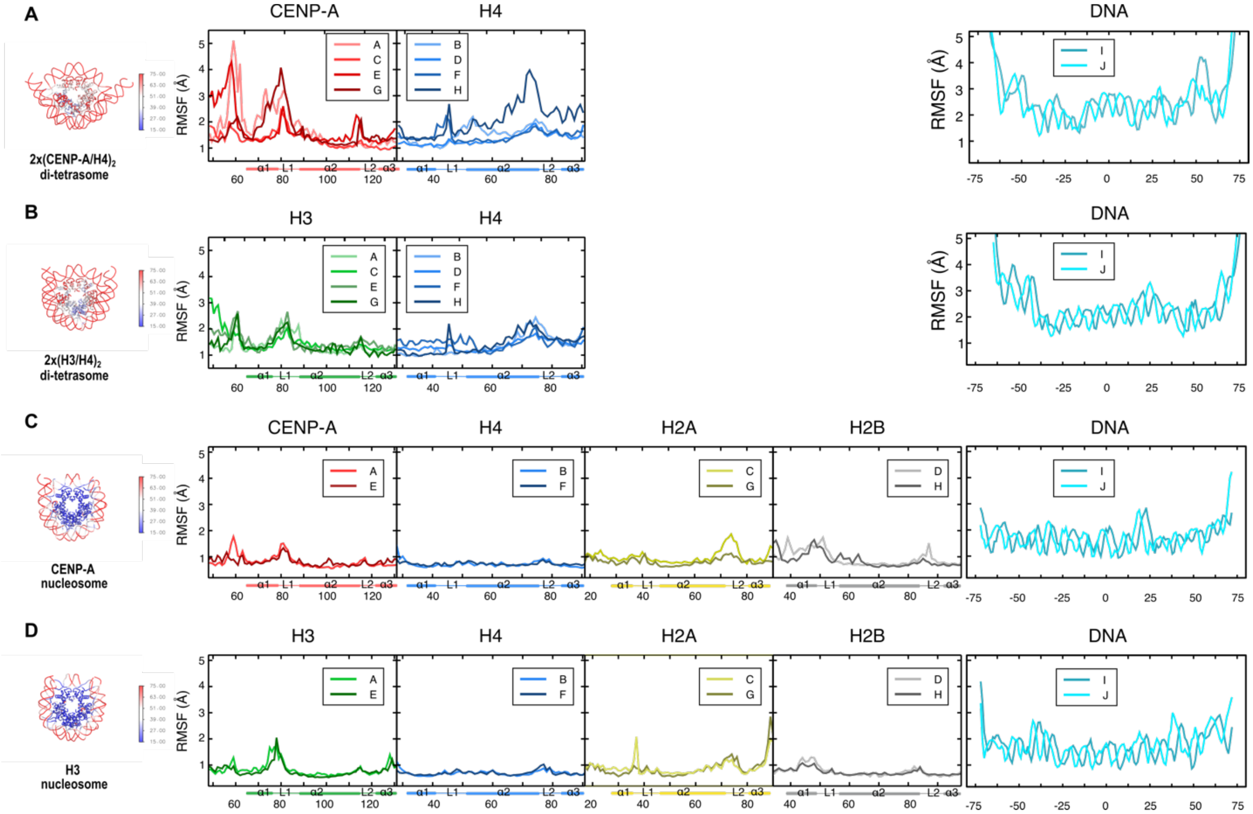


**Figure S6. Root-mean-square-fluctuations (RMSF) comparison of protein and DNA chains in di-tetrasomes and nucleosomes during MD simulations. A.** 2x(CENP-A/H4)$_2$ di-tetrasome. Note that 4 chains of CENP-A and 4 chains of H4 histone are present in the structure. The (CENP-A/H4)$_2$ dimers closer to the dyad are represented with chains C/D and E/F, while those closer to the DNA entry/exit site are A/B and G/H. **B.** 2x(H3/H4)$_2$ di-tetrasome. **C.** CENP-A nucleosome. **D.** H3 nucleosome. Due to the extremely high mobility of the histone tails, only the histone folding domains (HFD) are shown. The RMSF values for each system are plotted on the pdb structure on the left.

**A CENP-A nucleosome binds CENP-C$^{CR}$ using both electrostatic and hydrophobic patches**

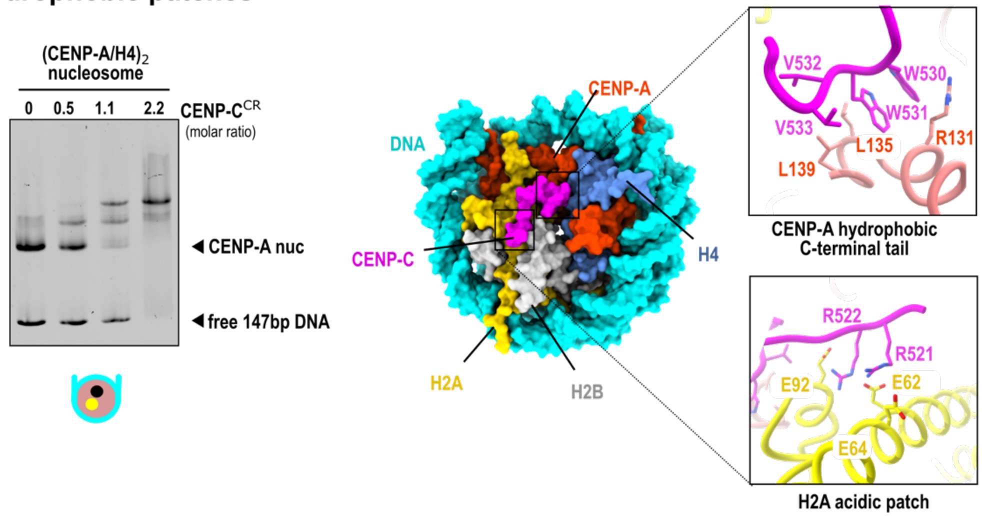


**B (CENP-A/H4)$_2$ tetrasome and di-tetrasome bind CENP-N$^{1-240}$**

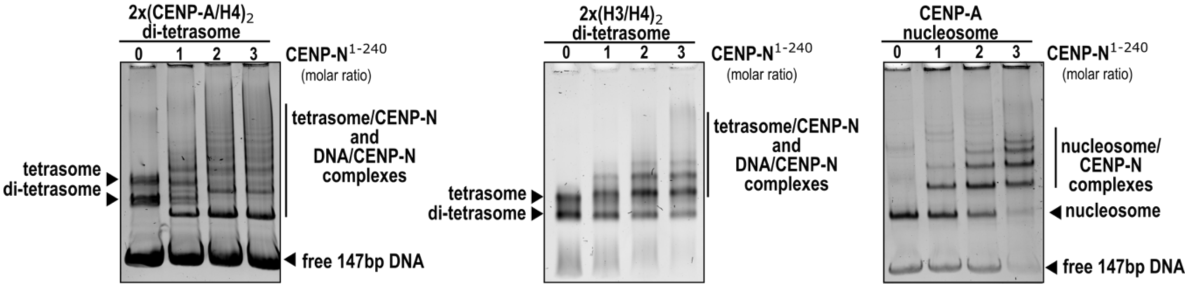


**Figure S7. A.** 5% native gel showing the binding of CENP-C$^{CR}$ to the CENP-A nucleosome (left). Note the appearance of a strong and defined band on the native gel after CENP-C$^{CR}$ binding, indicating a homogeneous and stable complex. A schematic of the CENP-A nucleosome is shown below the native gel with the hydrophic C-terminal tail of CENP-A (black circle) and the acidic patch of H2A (yellow circle) (only on one side of the structure). The histone core is red and the DNA is cyan. Note that unlike the tetrasome and di-tetrasome, the CENP-A nucleosome has two binding sites for CENP-C$^{CR}$ on each side, consistent with the increased binding affinity. (Right) Surface representation of the structure of the CENP-A nucleosome/CENP-CCR complex (PDB 6SE6) highlighting the two binding sites (hydrophobic C-terminal tail of CENP-A and H2A acid patch). CENP-A, H4, H2A, H2B, DNA and CENP-C are shown in red, blue, yellow, grey, cyan and magenta, respectively. The key residues responsible for the interaction with CENP-C$^{CR}$ in both binding sites are shown in sticks. **B.** 5% native gel showing the binding of CENP-N$^{1-240}$ to the (CENP-A/H4)$_2$ di-tetrasome. Note that the di-tetrasome band disappears immediately and is replaced by a lower band that likely corresponds to the (CENP-A/H4)$_2$ di-tetrasome-CENP-N complex. (Middle) Native gel showing the binding of CENP-N to the (H3/H4)$_2$ di-tetrasome. Note that the band corresponding to the (H3/H4)$_2$ tetrasome disappears faster than the band corresponding to the (H3/H4)2 di-tetrasome, indicating that CENP-N most likely binds to an unoccupied section of DNA in the (H3/H4)$_2$ tetrasome. (Right) Native gel showing the binding of CENP-N to the CENP-A nucleosome. Note that the binding shows the formation of larger complexes, indicating a polymerization process.

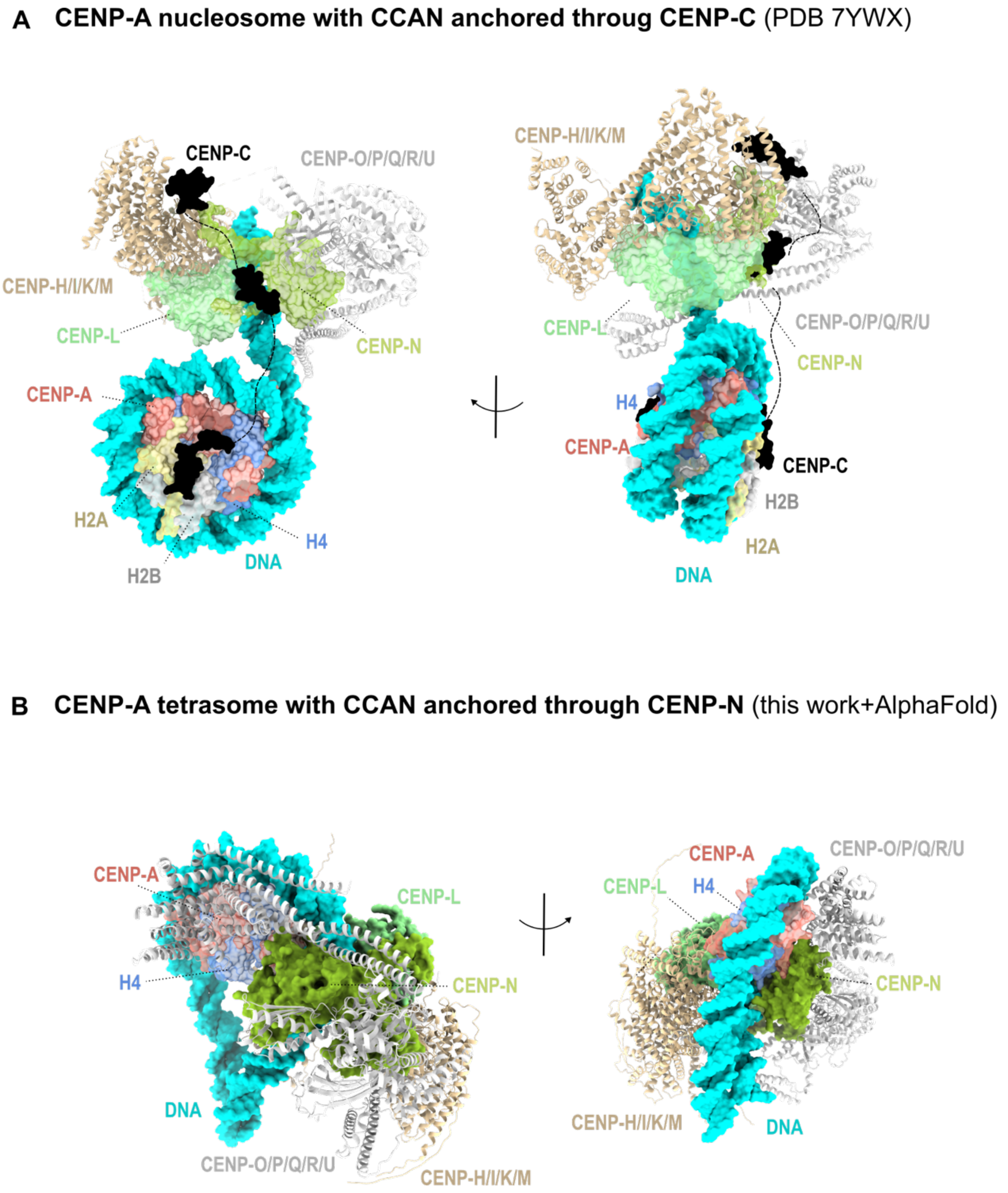


**Figure S8. A.** Space filling representation of CCAN bound to CENP-A nucleosome generated using PDB 7YWX **B.** Space filling representation of CCAN bound to CENP-A tetrasome generated using our structure of CENP-A tetrasome and AlphaFold modeling for CCAN complex. The structure was minimized and shows no steric clashes. CENP-A is red, H4 is blue, H2A is yellow, H2B is grey, DNA is cyan, CENP-C is black, CENP-N-L is green, CENP-H-I-K-M is beige, and CENP-O-P-Q-U-R is light gray. CENP-H-I-K-M and CENP-O-P-Q-U-R are shown as ribbon for clarity.
